# Serendipity and the slime mold: a visual survey of megadalton protein assemblies reveals the structure of the polyketide synthase Pks16

**DOI:** 10.1101/2025.03.12.642832

**Authors:** Gabriel Hoogerbrugge, Adrian T. Keatinge-Clay, Edward M. Marcotte

## Abstract

Large macromolecular assemblies are integral to most cellular processes, making their identification and structural characterization an important strategy for advancing our understanding of protein functions. In this pilot study, we investigated large multiprotein assemblies from the cytoplasm of the slime mold *Dictyostelium discoideum* using shotgun-electron microscopy (shotgun-EM), the combined application of mass spectrometry-based proteomics and cryo-electron microscopy (cryo-EM) to heterogenous mixtures of proteins. With its similarities in cell structure and behavior to mammalian cells, *D. discoideum* has long served as an invaluable model organism, particularly in the study of immune cell chemotaxis, phagocytosis, bacterial infection, and other processes. We subjected *D. discoideum* soluble protein complexes to two-step fractionation, performing size-exclusion chromatography followed by mixed-bed ion-exchange chromatography. Isolated fractions containing a subset of megadalton-scale protein assemblies were subsequently analyzed using mass spectrometry to identify the proteins and cryo-EM to characterize their structures. Mass spectrometry analysis revealed 299 unique proteins in the isolated fractions, then single-particle cryo-EM analysis generated distinct 2D projections of several visually distinctive protein assemblies, from which we successfully identified and reconstructed three major protein complexes: the 20S proteasome, the dihydrolipoyllysine-residue succinyltransferase (Odo2) of the mitochondrial 2-oxoglutarate dehydrogenase complex, and polyketide synthase 16 (Pks16), thought to be the primary fatty acid synthase of *D. discoideum*. Based on the Pks16 structure, the first of the 40 *D. discoideum* PKSs to be experimentally determined, models for the full set of *D. discoideum* PKSs were constructed with help from AlphaFold 3. Comparative analysis enabled structural characterization of their reaction chambers. Shotgun-EM thus provides a view of proteins in their native or near-native biological conformations and scaling up this approach offers an effective route to characterize new structures of multi-protein assemblies directly from complex samples.

## Introduction

*Dictyostelium discoideum*, a soil-dwelling social amoeba commonly referred to as slime mold, has played a pivotal role in advancing our understanding of essential biological processes, including bacterial infection, immune cell chemotaxis, phagocytosis, and mitochondrial and neurological disorders. *D. discoideum* is an excellent organism for investigating diverse aspects of eukaryotic cell biology, as it exhibits structural and behavioral characteristics akin to mammalian cells and shares conserved intracellular signaling pathways (1), while also having rich repertories of genes relevant to its alternate unicellular/multicellular lifestyle and bacterial predation activities. Due to its extensive repertoire of conserved functions, *D. discoideum* also serves as an accessible system for investigating human protein orthologs, situated within a complexity range intermediate between *S. cerevisiae* and higher multicellular eukaryotes (1).

Of particular note, *D. discoideum* contains a rich repertoire of genes encoding polyketide synthases (PKSs), highlighting a potentially active secondary metabolic network involved in the biosynthesis and export of small molecules (2). PKSs are responsible for the synthesis of polyketides, which can have antibiotic and signaling activities. These compounds have significant applications in the pharmaceutical, agrochemical, and biotechnological industries (3, 4). Although a wide range of PKS genes are present in eukaryotic organisms, the ecological functions of the resulting polyketides—beyond their pharmaceutical utility—remain widely underexplored. Understanding the ecological roles of polyketides is essential for advancing our comprehension of microbial cellular processes and their interactions within the environment (5). *D. discoideum* has approximately 40 putative PKS genes (2), but the functional roles and structural characteristics of these PKSs are poorly understood, as only a limited number of their products have been isolated and characterized, and no structures of *Dictyostelium* PKSs have been solved to date.

There are numerous advanced techniques for isolating specific proteins of interest, including recombinant expression and affinity- or immuno-purification. However, methods that rely on protein tagging or antibodies have certain drawbacks. Specifically, the biological system must be receptive to genetic manipulation, and the addition of tags may destabilize protein-protein interactions (6) or alter protein stoichiometry, while immunopurifications require available antibodies, often lacking in less studied species. In contrast, structural analysis of endogenous protein complexes offers a promising alternative, as it bypasses the challenges posed by recombinant expression (7) and tagging techniques. Such “tagless” approaches to isolating protein assemblies can permit the unbiased analysis of cellular lysates, providing a more natural representation of protein interactions (6).

One such “tagless” approach involves directly examining endogenous protein complexes using cryo-EM (7–10), which has revolutionized the structural characterization of large, flexible, and heterogeneous biomolecular complexes. The “resolution revolution,” driven by the advent of direct detectors that enhance the signal-to-noise ratio in electron micrographs, along with advancements in automation for data acquisition and image processing, has enabled cryo-EM to be applied effectively for studying molecular organization in both highly purified isolated biomolecules and *in situ* (11, 12). However, it is still challenging to apply cryo-EM to heterogenous mixtures, as increased sample complexity complicates identifying individual assemblies and assigning individual particles to their corresponding structures, and the corresponding lower particle counts can lead to lower-resolution reconstructions.

Nonetheless, an emerging field within structural proteomics focuses on developing and improving methods to investigate molecular structures directly within native cell extracts. In particular, mass spectrometry (MS) provides a powerful complementary approach to determine the molecular composition of the samples. Its high sensitivity, specificity, and ability to generate both structural and quantitative data make MS particularly valuable when integrated with cryo-EM. This combined approach, sometimes referred to as shotgun cryo-EM (9), allows for the acquisition of high-resolution structures of multiple, unrelated protein complexes from a single cryo-EM dataset (7, 13). When applied to crude extracts of intermediate complexity, this approach can enhance *in situ* techniques, such as cryo-focused ion beam tomography, by directly correlating structural data with molecular signatures seen in tomograms.

This study represents a pilot investigation of employing the shotgun cryo-EM approach on *D. discoideum* cell lysates in order to characterize several of its megadalton multiprotein assemblies. We combined mass spectrometric analysis for protein identification and cryo-EM for structural characterization on macromolecular size via size-exclusion chromatography (SEC), followed by separation based on surface charge using mixed-bed ion-exchange chromatography (IEX). Using this integrated approach, we reconstructed four distinct protein complexes, including one yet to be identified. Notably, we serendipitously solved a 3.9 Å structure of the first *D. discoideum* PKS, Pks16. Pks16 is thought to be the primary fatty acid synthase (FAS) of *D. discoideum* (14). Hence, the structure provides an initial experimental view of this important protein family in *D. discoideum*, enabling a comparative structural analysis across the family members.

## Experimental procedures

### Experimental design and statistical rationale

A series of biological replicates were used to establish reproducible chromatographic separation and enrichment procedures. For the sample ultimately analyzed by cryo-EM (a pool of 3 HPLC fractions), protein mass spectrometry analysis of the identical sample was performed as a technical triplicate experiment, and confirmed with analysis of the 3 corresponding fractions from a biological replicate. Mass spectrometry proteomics data were analyzed with Proteome Discoverer (ThermoFisher) as detailed below, controlling protein and peptide identification false discovery rates at <1% FDR, and are reported in **Table S1**. Cryo-EM statistics were computed with cryoSPARC v4.5 and are detailed in full in **Table S2** and **Supplemental Figures S1-S4**, including micrograph and particle counts, 3D reconstruction processing statistics (including FSC thresholds and computed map resolutions), and model refinement and validation statistics as calculated by Phenix (15). 3D structural models were computed initially with AlphaFold 3 (16) before refining subsequently with Namdinator (17).

### Dictyostelium strain and culture

*Dictyostelium discoideum* AX2-214 cells, obtained from the Dicty Stock Center (DSC) at Columbia University, New York, were cultured in HL5 axenic medium (Formedium), supplemented with 74.8 mM glucose, 10,000 units/mL of penicillin, and 10,000 µg/mL of streptomycin. Cultures were maintained at 22°C in cell culture plates, with shaking at 180 rpm in culture flasks.

### Dictyostelium lysis and soluble protein extraction

*D. discoideum* cultures were grown in 3 L of medium to a cell density not exceeding 4×10^6^ cells/mL. Cells were harvested by centrifugation at 500 × g for 4 minutes at room temperature. The resulting cell pellets were resuspended in 400 mL of 50 mM HEPES, pH 7.4, repelleted, and then suspended in 8 mL of Dicty Lysis Buffer (50 mM HEPES, pH 7.4, 100 mM NaCl, 3 mM MgSO_4_, 0.1 mM EGTA, 1 mM DTT, 0.1 mM PMSF, cOmplete mini EDTA-free protease inhibitors (Roche), PhosSTOP EASY phosphatase inhibitors (Roche), and 1% Igepal CA-630 (Sigma-Aldrich)). Cells were lysed by incubation on ice with vortexing every minute. To degrade nucleic acids, benzonuclease was added to a final concentration of 1 unit/mL and incubated on ice for 30 minutes. Following incubation, the lysate was centrifuged at 17,000 × g for 20 minutes at 4 °C to remove intact cells, organelles, and insoluble proteins. The supernatant was then subjected to ultracentrifugation at 100,000 × g for 1.25 hours at 4 °C to remove membrane fragments and residual debris. The protein concentration in the detergent-solubilized cytosolic fraction was quantified using a DC Bradford Assay (BioRad).

### Dictyostelium soluble protein separation

Two milliliters of cytosolic soluble extract in HEPES-Dicty Lysis Buffer were fractionated by size exclusion chromatography with a preparative-grade HiLoad 16/600 Superdex 200 PG column (Cytiva) at a flow rate of 1 mL/min. The mobile phase consisted of 50 mM HEPES, pH 7.4, 100 mM NaCl, and 3 mM MgSO_4_, 0.1 mM EGTA. Fractions of 1.5 mL were collected from the megadalton region, pooled, and concentrated using a Sartorius Vivaspin Turbo 100,000 MWCO ultrafiltration unit. The concentrated pooled fractions from the megadalton region were further fractionated using a mixed-bed ion exchange column (PolyLC Inc, #204CTWX0510) with a Dionex UltiMate 3000 HPLC system. Chromatographic separation was achieved by applying a gradient of Buffer A (50 mM HEPES, pH 7.4, 3 mM MgSO_4_, 0.1 mM EGTA) and Buffer B (50 mM HEPES, pH 7.4, 1.5 M NaCl, 3 mM MgSO_4_, 0.1 mM EGTA). Fractions of 500 µL were collected into a 96-well plate and subsequently analyzed by mass spectrometry.

### Protein mass spectrometry

Proteins were identified and quantified using a Thermo Orbitrap Fusion Lumos tribrid mass spectrometer. Peptide separation was performed via reverse-phase chromatography on a Dionex Ultimate 3000 RSLCnano UHPLC system (Thermo Scientific), employing a C18 trap coupled to an Acclaim C18 PepMap RSLC column (Dionex; Thermo Scientific). Mass spectra were acquired using a standard top-speed HCD MS1-MS2 method, and the data were analyzed using the Proteome Discoverer standard workflow (ThermoFisher).

Mass spectra files were analyzed using Proteome Discoverer 2.3. The spectra were searched against the complete *Dictyostelium discoideum* proteome obtained from UniProt, as well as a contamination database provided by the Hao group (18). Tryptic peptides were considered with a maximum of two missed cleavages. The analysis was performed with a minimum precursor mass of 350 Da and a maximum precursor mass of 6,000 Da, applying a precursor mass tolerance of 10 ppm. FDR targets for peptide identification were set with a strict threshold of <1%. The FDR criterion for protein identification mirrored those for peptide identification, with a strict target of <1% and a relaxed target of <5%. The proteins listed in **Table S1** represent only those with high-confidence identifications, corresponding to an FDR of <1%, with peptide identifications provided in the supporting tab of **Table S1**.

### Cryo-EM sample preparation and data collection

Concentrated protein fractions from the mixed-bed ion exchange chromatography were adjusted to a final composition of 50 mM HEPES, pH 7.4, 100 mM NaCl, 3 mM MgSO4, 0.1 mM EGTA, 1 mM DTT, 0.1 mM PMSF, Roche cOmplete mini EDTA-free protease inhibitors, Roche PhosSTOP EASY phosphatase inhibitors, and 2% glycerol. This adjustment was made to lower the salt concentration and incorporate a cryoprotectant for subsequent plunge-freezing. A 3 µL aliquot of the concentrated protein solution was applied to a glow-discharged C-Flat Holey Thick Carbon 1.2/1.3 Cu400 grid, maintained at 100% humidity at 4 °C. Using an FEI Vitrobot Mark IV (Thermo Fisher Scientific), the grid was blotted for 10 s with a blot force of 1 and a wait time of 5 s, before being plunged into liquid ethane. A total of 332 micrographs were collected using a 200 kV FEI Glacios (Thermo Fisher Scientific) cryo-transmission electron microscope equipped with a Falcon IV direct electron detector (FEI). Exposures were collected with a calibrated pixel size of 0.933 Å/pixel, a total electron dose of 49 e−/Å², and a defocus range between −1.5 and −2.5 µm.

### Cryo-EM data processing, building, and refinement

On-the-fly data processing was conducted using cryoSPARC Live (19) encompassing motion correction, defocus estimation, micrograph curation, and particle picking. Subsequent data processing steps, including particle extraction, curation, and refinement, were performed using cryoSPARC v4.5 (20). The EM processing pipelines for the four reconstructed complexes are presented in full in **Figures S1-S4**, and are summarized in brief as follows.

For the 20S proteasome complex, particles selected from 2D classification were subjected to an *ab initio* 3D reconstruction followed by homogeneous refinement, with no symmetry applied (C1). This was followed by a second homogeneous refinement with C2 symmetry imposed, culminating in a final non-uniform refinement, again with C2 symmetry.

For the Odo2 complex, multiple rounds of 2D classification and *ab initio* reconstruction were carried out to refine the particle stack. Selected particles underwent an *ab initio* 3D reconstruction, followed by homogeneous refinement (C1). A subsequent homogeneous refinement with octahedral (O) symmetry was performed, and duplicate particles were removed prior to a final homogeneous refinement with O symmetry imposed.

For the hexameric star complex, multiple rounds of 2D classification and ab initio reconstruction were employed to refine the particle stack. After selecting the best class from 2D classification, an *ab initio* 3D reconstruction was followed by homogeneous refinement without symmetry (C1), and a final non-uniform refinement was performed with no symmetry imposed (C1).

For Pks16, iterative 2D classification and *ab initio* reconstruction were performed to refine the particle stack. Particles selected from 2D classification underwent *ab initio* 3D reconstruction, followed by homogeneous and non-uniform refinement on the best class with C2 symmetry. The particle stack was further refined by removing duplicate particles, with final homogeneous and non-uniform refinements performed with C2 symmetry. Templates were generated from the volume obtained in the previous non-uniform refinement. A template picker was then employed to select particles, which underwent *ab initio* reconstruction followed by homogeneous and non-uniform refinement, again with C2 symmetry imposed. The final particle stack was refined by removing duplicate particles, with the final homogeneous and non-uniform refinements conducted with C2 symmetry.

Initial structural models were generated with AlphaFold 3 and visualized using UCSF ChimeraX (21). The N-terminal domain was independently fit to density using Isolde (22), the model optimized by Namdinator, and the full model refined in Phenix.

### AlphaFold 3 modeling of DdPKSs

AlphaFold 3 correctly predicted the structure of Pks16, so we asked it to predict the structures of the KS-ACP regions for the other 39 *Dd*PKSs. PKSs were split into 2 parts, as 4920 residues is the limit with 4 NADPH molecules. To preserve homodimeric interactions, KS+AT+DH and KS*+DH+MT+KRs+ER+KRc+ACP dimers were independently predicted (KSATDH_PksXX.pdb and noAT_PksXX.pdb Data Files; KR_s_ and KR_c_ are the structural and catalytic subdomains of KR) and superposed through the KS dimer (* indicates 3 KS residues at Pks16 positions 86-88 were changed to aspartate to avoid the dominant KS/ACP association, **Figure S5**). As the merged models contain KS+AT from the first structure and DH+MT+KRs+ER+KRc+ACP from the second structure, they also contain a small defect at their junction, downstream of the conserved proline at position 935 (Merged_PksXX.pdb Data Files). NADPH is equivalently bound to each KR but predicted to bind through diverse modalities to ER and not predicted to bind Pks10 ER. The prediction for the unusual KS+AT+DH+KRs+ER+KRc+ACP portion of Pks37 was obtained by supplying AlphaFold 3 with 2 copies of residues 4-2503 from Pks37.

To investigate the association of KS and ACP, 2 copies of KS+AT and ACP were supplied to AlphaFold 3 for each *Dd*PKS. A consensus association was identified (KSATACP_PksXX.pdb Data Files); it is similar to the KS/ACP association formed during the extension reaction in modular PKSs (23, 24). To investigate the association of AT and ACP, one copy of both the AT and ACP domain was supplied. From the 200 returned predictions (5 per job), a consensus docking site was identified (ATACP_PksXX.pdb Data Files); it is similar to the AT/ACP association in modular PKSs (25). To investigate the association of MT and ACP, a consensus docking site was identified from the KS*+DH+MT+KRs+ER+KRc+ACP dimers, and more examples were obtained through supplying MT+KR+ACP monomers to AlphaFold 3 (MTACP_PksXX.pdb Data Files). To investigate the association of KR and ACP domains, a consensus docking site was identified from the KS*+DH+MT+KRs+ER+KRc+ACP dimers, and more examples were obtained through supplying MT+KR+ACP monomers to AlphaFold 3 (KRACP_PksXX.pdb Data Files). To investigate the association of DH and ACP domains, a consensus docking site was identified from the KS*+DH+MT+KRs+ER+KRc+ACP dimers (DHACP_PksXX.pdb Data Files). No consensus docking site was observed from isolated DH and ACP domains. To investigate the association of ER and ACP domains, a consensus docking site was identified from the KS*+DH+MT+KRs+ER+KRc+ACP dimers (ERACP_PksXX.pdb Data Files); it is similar but distinct from the ER/ACP association in modular PKSs (26). Structures containing consensus associations were superposed with corresponding Pks16 domain(s) for analysis (for KS/ACP, the KS dimer; for AT/ACP, the AT of chain B; for MT/ACP, the large MT subdomain of chain A; for KR/ACP, the KRs of chain A; for DH/ACP, the DH of chain A; for ER/ACP, the ER of chain A).

## Results and discussion

### Fractionation and enrichment for megadalton macromolecular assemblies

We sought to apply shotgun-EM to survey high abundance megadalton protein complexes present in *Dictyostelium* slime mold cell lysate. As the complexity of unfractionated cell lysate is too high to solve individual structures, we first performed chromatography to reduce the complexity. In this study, we employed a rapid and efficient two-step biochemical fractionation of detergent-solubilized lysates from *D. discoideum* to isolate large macromolecular assemblies in a near-native state. Prior to fractionation, the protein assemblies were ultracentrifuged to prevent liposome formation in downstream processes. Native protein complexes were then separated according to molecular mass using size-exclusion chromatography. SEC has become a widely adopted fractionation technique (7, 9), with the selection of high-molecular-weight fractions offering the advantage that larger proteins are more visually evident by transmission electron microscopy. Three high molecular weight fractions were collected from SEC and subjected to mixed-bed ion-exchange chromatography to further separate the macromolecular assemblies based on surface charge properties. Three IEX fractions were then analyzed in parallel by mass spectrometry for proteomic identification and by cryo-EM for structural characterization of the assemblies (**Figure 1A**). As this was an initial pilot study to evaluate feasibility of this method for future scale-up, we have not yet exhaustively analyzed the remaining fractions.

**Figure 1.**
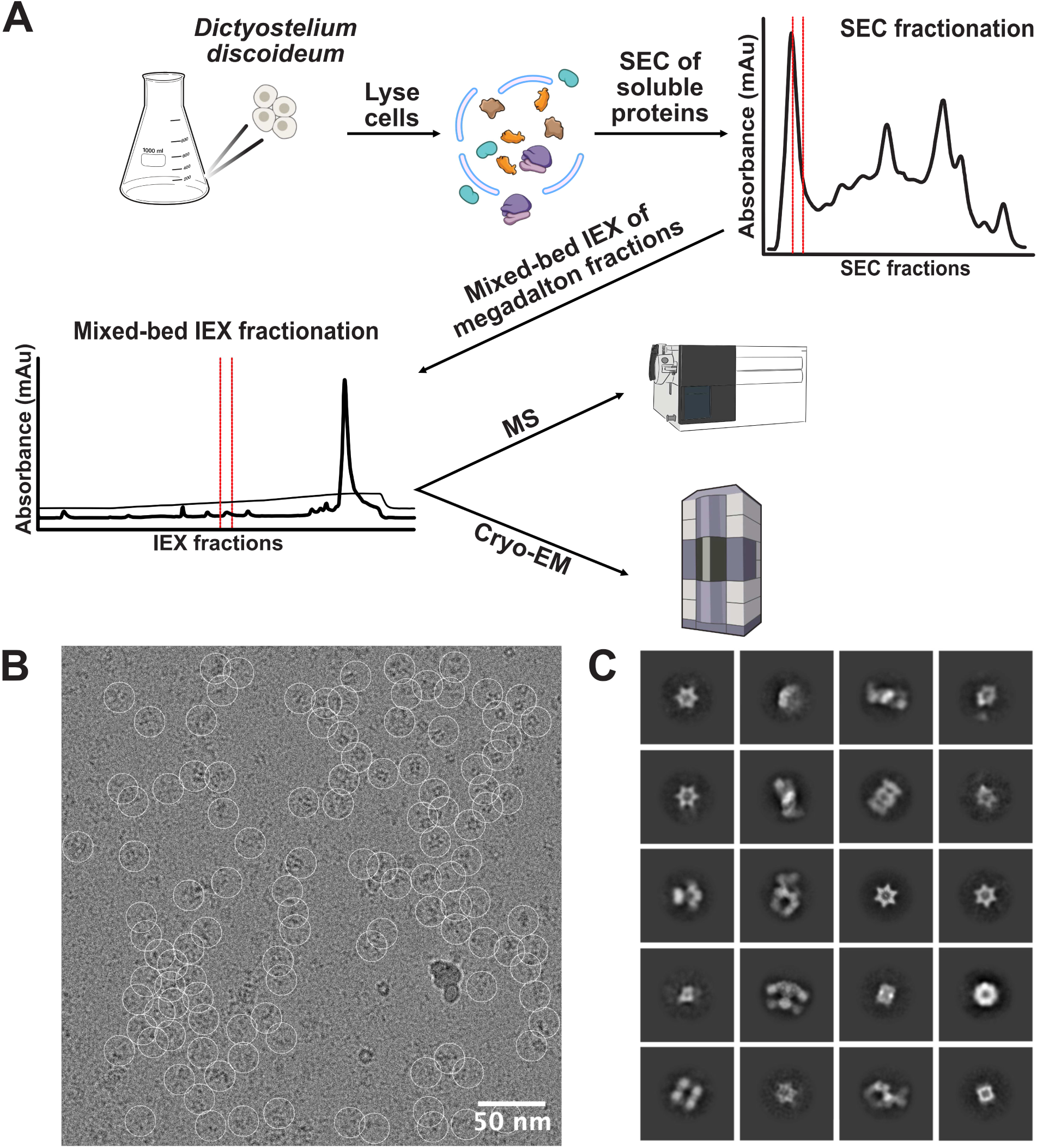
Overview of a small pilot survey of megadalton complexes in *Dictyostelium discoideum.* (A) Cells were collected during their vegetative stage and lysed using Igepal. The detergent soluble cytosolic proteins were first fractionated using SEC chromatography then by mixed-bed IEX to further reduce the heterogeneity of the protein mixture. Proteins enriched from SEC-IEX fractionation were loaded on cryo-EM grids and visualized, and in parallel, proteins were identified by MS. (B) Cryo-EM micrograph of proteins from the concentrated mixed-bed IEX fraction. Particles were picked using cryoSPARC live using template-free auto-picking (white circles). Scale bar, 50 nm. (C) 2D class averages for a subset of protein complexes with distinctive shapes. Class averages were computed using a 24 nm diameter circular mask.

In particular, cryo-EM enables the reconstruction of densities for macromolecular assemblies, but associating protein identities with these densities can be challenging without a comprehensive catalog. To facilitate this process, we employed MS in parallel, enhancing the identification and assignment of protein complexes to their corresponding EM densities. Following mixed-bed IEX, the fractions were subjected to trypsin digestion, and the resulting tryptic peptides were analyzed by liquid chromatography-tandem mass spectrometry (LC-MS/MS). MS analysis identified 239 unique proteins, with 125 exhibiting six or more peptide spectral matches (PSMs) (**Table S1**). This catalog of proteins captured a broad range of cellular and molecular functions, which guided and validated the identification and reconstruction of macroassemblies.

### Cryo-EM, 2D classification, and single-particle analysis of enriched complexes

To investigate the structural characteristics of the isolated protein complexes, cryo-EM was employed on the fractions obtained following IEX. A total of 332 micrographs were collected over a three-hour period. Real-time processing was conducted using cryoSPARC Live, which included motion correction, defocus estimation, and particle selection. Template-free auto-picking facilitated the identification of over 37,000 particles (**Figure 1B**). Subsequent 2D classification of these particles generated projections representing various orientations of the protein complexes, revealing several visually distinct structures (**Figure 1C**). Notably, recognizable complexes such as the 20S proteasome, which is readily identifiable in complex mixtures (7–9), were observed.

A key challenge in cryo-EM analysis of heterogeneous mixtures is the identification of enriched protein complexes and the accurate assignment of corresponding densities to specific protein assemblies (14). By reducing sample complexity, we significantly improved the ability to match multiple 2D projections from the same complexes. The resulting 2D classes exhibit distinct features that enable effective sorting for further processing without exhaustive combination testing. For instance, several of the 2D classes display a hexameric star-like formation, which is clearly distinguishable from the well-characterized heptameric proteasome, allowing for the straightforward identification and grouping of all 2D projections exhibiting a size-matched hexameric arrangement.

After sorting the 2D classes, *de novo* reconstructions were generated and utilized in conjunction with the MS-derived identities, providing a foundation for subsequent refinement. The application of symmetry during reconstruction necessitates an understanding of the EM density to prevent the introduction of bias or artifacts that could undermine the resolution of the reconstructed structure. With the protein complexes identified through MS analysis, we were able to more accurately associate 2D projections with their corresponding protein assemblies and apply symmetry constraints appropriately, leading to improved structural models.

### Identification of candidate complexes by comparing mass spectrometry and cryo-EM

One of the most prominent 2D classes identified was the proteasome, with MS data detecting all 14 subunits (7 α and 7 β subunits) of the 20S proteasome (**Figure 2A**). Additionally, we observed a 2D class displaying a quadrilateral, box-like arrangement with four distinct corners, which corresponded to an EM density exhibiting octahedral symmetry. Enforcing O-symmetry brought the reconstruction from 12.04 Å (C1) to 5.72 Å (O) resolution, considerably improving the quality of the cryo-EM density map. Based on the density, we could identify the protein as Odo2, the dihydrolipoyllysine-residue succinyltransferase subunit of the mitochondrial 2-oxoglutarate dehydrogenase complex, which forms a complex with octahedral symmetry and was present at high abundance in the mass spectrometry data (**Figure 2B**).

**Figure 2.**
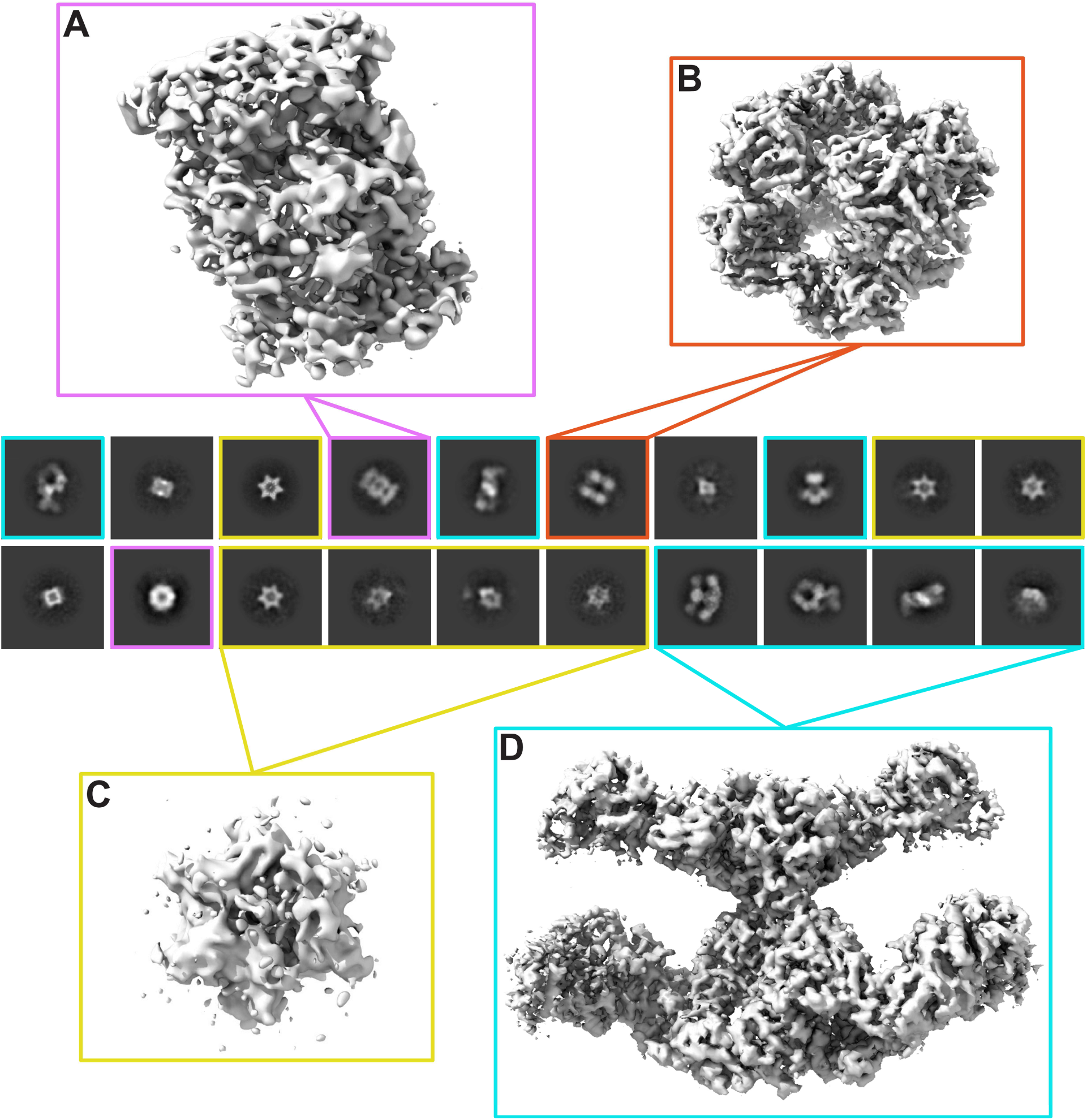
Cryo-EM 2D-class average projections and electron densities of the major protein complexes observed. Class average projections are outlined in colored squares by the corresponding protein complex: (A) 20S proteasome, purple; (B) dihydrolipoyllysine-residue succinyltransferase component of mitochondrial 2-oxoglutarate dehydrogenase complex, orange; (C) unidentified 6-pointed star-shaped complex, yellow; and (D) polyketide synthase 16, light blue.

Among the other particles, we observed a distinct hexameric star-like structure that did not obviously correspond to any of the high-abundance, well-characterized proteins in the sample (**Figure 2C**), as well as an extended structure that was more readily identified as the most abundant protein in the sample, polyketide synthase 16 (Pks16), as indicated by both cryo-EM particle count and MS, with a particle count of 5,611 and 3,222 total PSMs (**Tables S1, S2, Figure 2D**).

To identify the star-shaped macromolecular complex, we tested each high-abundance protein by modeling the protein with AlphaFold 3 (including considering multimeric assemblies) and comparing the models with the cryo-EM density. Two potential candidates emerged: (i) an ATP-dependent protease subunit HsLV protein with homology to known dodecameric *Escherichia coli* ATP-dependent protease subunit HsLV (PDB entry: 1E94) (27), or (ii) a partial assembly of the F1 ATP synthase. We first suspected the star might be a hexameric heat shock protein, and while hexameric heat shock protein HspA was highly abundant in the MS data, its predicted structure was a poor match for the Cryo-EM density. In contrast, a dodecameric assembly of the ATP-dependent protease protein HsLV protein fitted the density well. However, our MS data suggested that it was only present at very low abundance across the biological replicate experiments, which considerably reduced our confidence in this identification.

Thus, given the abundant visual representation of the hexameric structure in micrographs and particle counts from the 2D classes, we also considered ATP synthase as a potential candidate. ATP synthase was a plausible candidate, given its high abundance in the MS analysis and the reasonable alignment of the AlphaFold 3 model with the cryo-EM density. To this end, we used AlphaFold 3 to predict the structure of ATP synthase, incorporating three α-subunits, three β-subunits, and one γ-subunit to represent its natural assembly. The predicted model was then superimposed onto the low-resolution cryo-EM map density (**Figure 3A**). Visualization of the model within the cryo-EM map supports ATP synthase as a strong candidate, as the predicted structure aligns with the EM density and retains key features, such as the density in the central cavity, where the γ-subunit is expected to reside. In contrast, the ATP-dependent protease subunit HslV protein complex, while exhibiting a well-defined hexameric structure when viewed down the central axis, does not account for the central density feature observed in the 2D classification, further distinguishing it from ATP synthase (**Figure 3B**).

**Figure 3.**
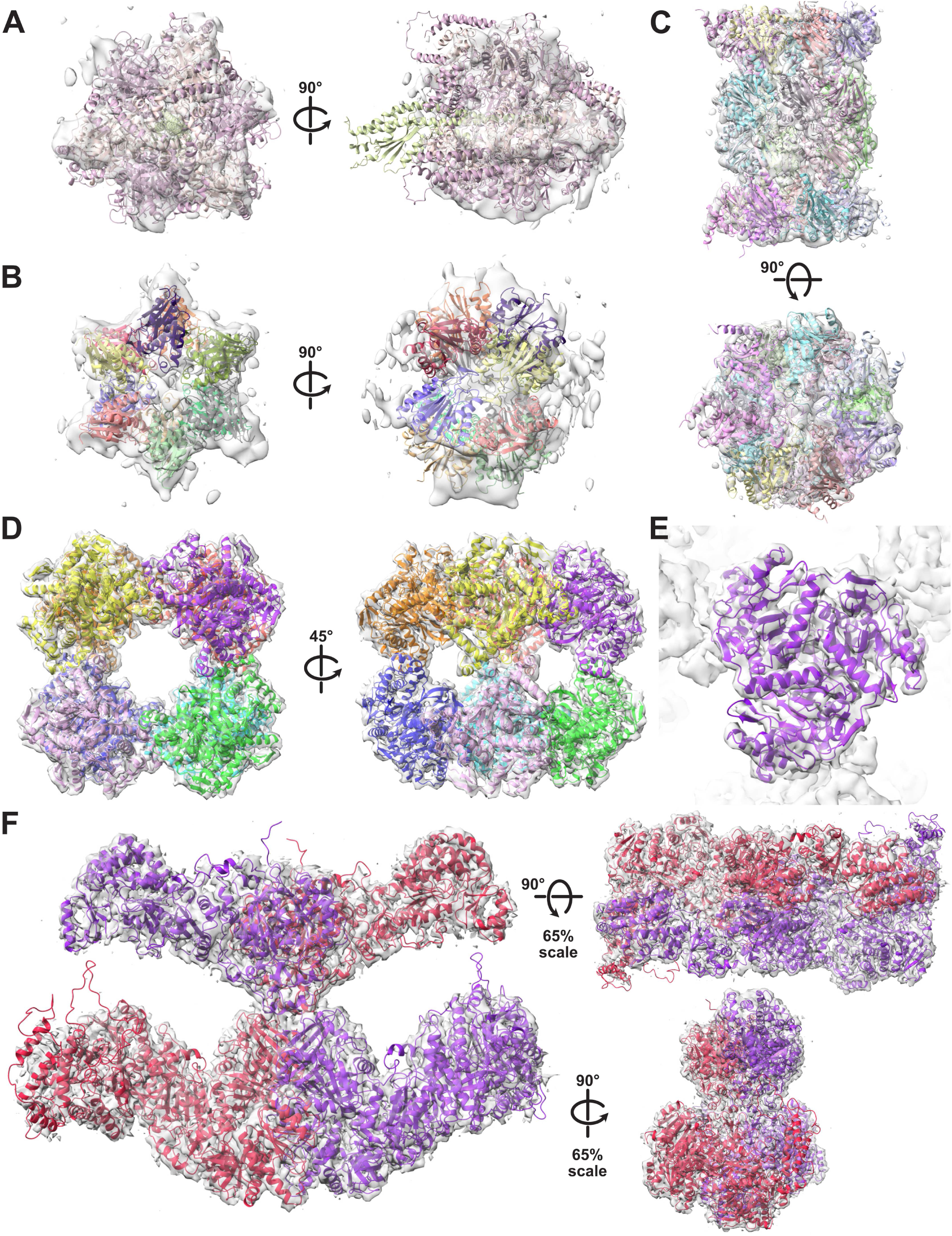
Molecular models of reconstructed protein complexes. (A) *D. discoideum* ATP synthase model, predicted using AlphaFold 3, docked into the refined low-resolution map density of the hexameric star-like assembly. (B) *D. discoideum* 12-mer structure of HsLV docked into the refined low-resolution map density of the hexameric star-like assembly. (C) *Homo sapiens* 20S proteasome (PDB 5LF1), serving as a proxy for the slime mold orthologous structure, docked into the refined 8.64 Å resolution map density. (D) *D. discoideum* Odo2 mitochondrial dihydrolipoyllysine-residue succinyltransferase subunit of the 2-oxoglutarate dehydrogenase complex, modeling residues 210-440 using AlphaFold 3, docked into the refined medium-resolution (5.89 Å) map density, with a magnified view of the trimeric arrangement in (E). (F) *D. discoideum* polyketide synthase 16, modeled using AlphaFold 3, docked into the refined 3.9 Å resolution map density.

However, MS only identified the β-subunit of ATP synthase (specifically, the AtpB protein), with no evidence for either the α- or γ-subunits. While there is literature support for homo-hexameric β rings in other classes of ATP synthase homologs (e.g. AAA+ ATPases (28), Rho (29)), they have not been documented for *D. discoideum* mitochondrial ATP synthase. Thus, we were not confident enough in either candidate (HsLV or AtpB) and this hexameric star structure remains unassigned.

### Modeling of reconstructed multiprotein assemblies

In order to build models for each of the complexes, we either computed candidate structures by AlphaFold 3 (**Figures 3A, B, D-F**) or alternatively, as a proxy for the *D. discoideum* 20S proteasome, we superimposed the *Homo sapiens* 20S proteasome experimental structure (PDB entry: 5LF1) (30) into our EM density (**Figure 3C**).

The dihydrolipoyllysine-residue succinyltransferase subunit of mitochondrial 2-oxoglutarate complex is previously known to form a homomeric 24-mer assembly exhibiting octahedral symmetry in *Homo sapiens* (31), so we modeled the *D. discoideum* Odo2 structure accordingly (**Figure 3D**). We employed AlphaFold 3 to predict a trimeric structure of Odo2 (amino acids 210-439) and docked into the cryoEM density map using ChimeraX. Despite the medium resolution (5.72 Å) of the map, the predicted model aligned extremely well with the features of the EM density (**Figure 3E**), especially in the sites of trimeric symmetry. Regions where AlphaFold 3 could not confidently predict structure (amino acids 1-209) were omitted, resulting in some voids in the density between the trimeric subunits, presumably occupied by the missing residues.

Finally, we predicted the structure of the *D. discoideum* Pks16 protein using AlphaFold 3 (amino acids 1-2496) and observed an excellent agreement overall in the cryo-EM density, further improved by subsequent computational refinement of the model into the density (**Figure 3F**). In general, the Pks16 density map exhibited higher resolution (<4.0 Å) in the core, while the outer arm and leg regions were of medium resolution (>6.0 Å), likely due to protein motion or the adoption of distinct conformations influenced by the presence of various molecules. Given the reduced resolution of these regions, we opted to use AlphaFold 3 for atomic model prediction rather than relying on alternative tools, such as ModelAngelo (32), for model generation directly in the cryo-EM density. The high consistency of the AlphaFold 3-predicted structure of the central body region, where ModelAngelo also generated a model, supported the use of AlphaFold 3 model for subsequent analyses and gave us confidence in constructing models for additional members of the *D. discoideum* PKS protein family.

### Modeling the full set of 40 Dictyostelium discoideum polyketide synthases

Pks16, thought to be the primary fatty acid synthase (FAS) of *D. discoideum* (Pks17 also operates during the late culmination stage) (14), is one of 40 highly-related, iterative type I PKSs in this social amoeba (2, 4, 33). As noted above, while we experimentally determined its structure by cryo-EM, we obtained essentially the same solution using the protein structure prediction tool AlphaFold 3. Since it cannot predict structures larger than 5000 residues, to better model dimeric interfaces, the 5206-residue Pks16 homodimer was predicted as 2 dimeric fragments and merged by superposing their KS domains (see **Experimental procedures**). Similar structures for 38 of the other PKSs were equivalently generated using AlphaFold 3 (Merged_PksXX.pdb Data Files, **Figures 4A & S5**). Because the functions of only a few of these PKSs are known, this represents an embarrassment of structural riches. From the 40 PKS structures, one can only hypothesize how Pks1 (also referred to as StlA) yields the short acyl chain incorporated into differentiation factor DIF-1 (34), how Pks5 yields the spore germination suppressing dictyodenes (5), and how Pks16 yields fatty acids (4) as the polyketides generated by the other PKSs have not been identified. While function lags structure for these PKSs, much can still be learned about how these multidomain machines operate through a comparative analysis of their structures.

**Figure 4.**
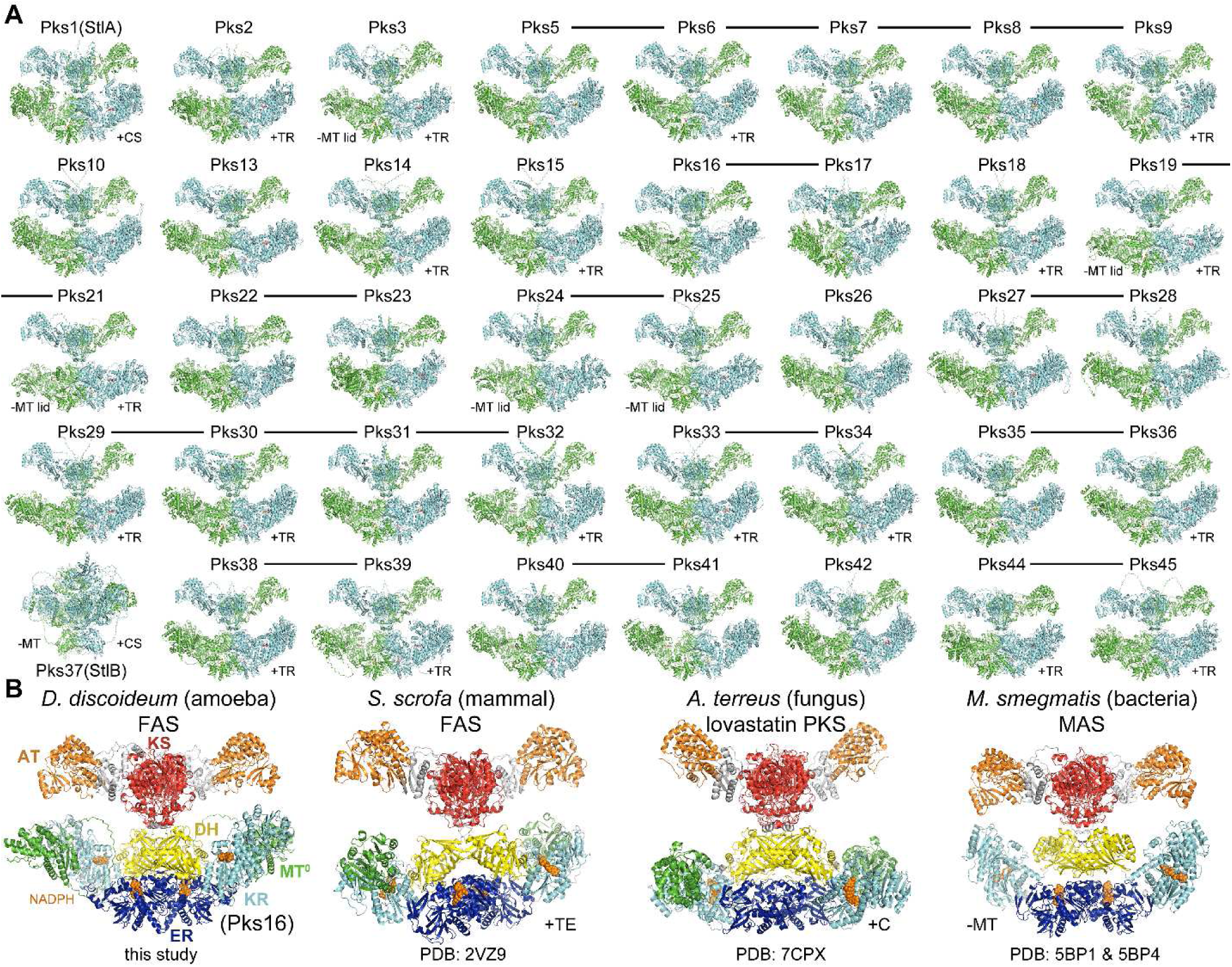
Structural relationships between Pks16 and other dimeric synthases from *D. discoideum* as well as mammalian, fungal, and bacterial species. (A) Pks16 is one of 40 *Dd*PKSs. With the exception of Pks37, the KS-ACP architectures predicted by AlphaFold 3 are highly similar (Merged_PksXX.pdb Data Files). Synthases whose genes are chromosomally adjacent and highly homologous are connected by lines (Ref. 4). Those possessing a C-terminal chalcone synthase (CS) or thioester reductase (TR) domain are indicated, as are those lacking an MT domain or MT lid (positions 1254-1414 and 1575-1594). (B) The principal *D. discoideum* FAS, Pks16, is compared with the porcine FAS, the lovastatin PKS, and MAS. The porcine FAS and lovastatin PKS possess a C-terminal thioesterase (TE) and condensation (C) domain, respectively. MAS does not contain an MT domain. PDB codes are indicated. AlphaFold 3 models for Pks16 and MAS are shown. KS, ketosynthase; AT, acyltransferase; DH, dehydratase; MT^0^, inactive methyltransferase; ER, enoylreductase; KR, ketoreductase.

The acyl carrier protein (ACP) domain plays a prominent role in these iterative type I PKSs, repeatedly docking to each of the other domains to perform the rounds of elongation and processing that yield the product(s) of a synthase (33). In the *Dd*PKSs, the other domains include an acyltransferase (AT) that transfers a malonyl extender unit from malonyl-CoA to the ∼18 Å-phosphopantetheinyl arm of ACP as well as a ketosynthase (KS) that acquires a processed intermediate from an acyl-ACP and fuses it with malonyl-ACP through a decarboxylative Claisen-like condensation to generate a β-ketoacyl-ACP intermediate. Processing domains may also be present, such as a methyltransferase (MT) that transfers a methyl group from *S*-adenosylmethionine (SAM) to the α-carbon of the β-ketoacyl-ACP intermediate, a ketoreductase (KR) that can set stereochemistries at the α- and β-positions as it uses NADPH to generate a β-hydroxyacyl-ACP intermediate, a dehydratase (DH) that eliminates water to generate an α/β-enoyl-ACP intermediate, an enoylreductase (ER) that can set stereochemistry at the α-position as it uses NADPH to reduce the α/β-enoyl-ACP intermediate, and an offloading enzyme that releases the product(s) from the synthase. While a processing enzyme may be present in a synthase, it may not be functional. Likewise, while a processing enzyme may be functional, it may not operate every round. In Pks5, DH does not work on intermediates smaller than a triketide, and MT does not work on intermediates smaller than a heptaketide. Processing enzymes may also perform different chemistry on different acyl-ACP intermediates, as in the hypothemycin PKS where KR generates an L-oriented hydroxyl group when reducing the diketide intermediate and a D-oriented hydroxyl group when reducing longer intermediates (35). Thus, the detailed interactions between acyl-ACPs and other domains within a PKS control the product(s) it generates.

### Synthase architecture

The iterative type I synthases responsible for biosynthesizing fatty acids for an organism (FASs) usually possess one of two very different domain organizations (36). A hexameric architecture is present in the FASs of fungi and mycolic acid-producing bacteria, whereas a dimeric architecture is present in the FASs of non-fungal, non-plant eukaryotes. While fungi and mycolic acid-producing bacteria employ hexameric synthases to generate fatty acids, they also utilize dimeric synthases to generate diverse non-fatty acid polyketides. As examples, the fungus *Aspergillus terreus* and the mycolic acid-producing *Mycobacterium smegmatis* respectively produce lovastatin and mycocerosic acid using PKSs with architectures equivalent to dimeric FASs. Organisms with dimeric FASs do not usually possess additional iterative type I PKSs; however, social amoeba are a marked exception. Distinguishing their FAS from their PKSs based on sequence is not trivial, and the structural information reported here does not aid much either, since Pks16 superposes well with 38 of the other 39 PKSs.

Pks16 can be compared to the few other dimeric synthases whose structures have been experimentally determined – the FAS from the mammal *Sus scrofa* (pig) (PDB: 2VZ9) (37), the lovastatin PKS from the fungus *A. terreus* (PDB: 7CPX) (38), and the mycocerosic acid synthase (MAS) from the bacterial species *M. smegmatis* (PDBs: 5BP1 & 5BP4) (**Figure 4B**) (39). In each of the synthases, monomeric AT domains are rigidly connected to a KS dimer through a flanking subdomain (FSD) (40), and monomeric KR domains are connected to DH and ER dimers that make contact with one another on the twofold axis. With the exception of MAS, a monomeric MT domain inserts into a loop of the KR structural subdomain, KR_s_. Whereas the KS+AT region is rigidly connected to the DH dimer in Pks16 and the lovastatin PKS, it is flexibly connected in the porcine FAS and MAS. ACP domains are not observed in any of these structures, nor are the monomeric offloading enzymes of the porcine FAS or the lovastatin synthase [a thioesterase (TE) and condensation domain, respectively] (41).

In the *Dd*PKSs, the tight interaction of 3 regions (positions 53-59, 926-937, and 959-962; position numbering is in reference to Pks16, **Figure S5**) unite the KS and DH domains (NoAT_PksXX.pdb Data Files). A major contact is made by a highly conserved proline at position 935 that inserts into a pocket formed by residues at positions 53, 55, 56, and 59. Another is from the side chain of a residue at position 937 (most commonly isoleucine) inserting into a pocket formed by residues 55, 58, and 59. Another is from the side chain of the residue at position 59 (most commonly a leucine) inserting into a pocket made by residues at positions 935-937 and 959-962. These interactions ensure that the PKS monomers in the X-shaped dimer are linearly organized. The KS/DH connection in the lovastatin PKS is similar yet distinct. Its interface is larger and proportionally more contact is made at its center.

In the *Dd*PKSs, the DH/KR interface is primarily mediated by complementary interactions between residues in the last β-strand of the DH β-sheet (positions 1073-1076) and a KR_c_ helix (positions 2272-2283) (NoAT_PksXX.pdb Data Files). Most commonly, a phenylalanine at position 1073 inserts into a pocket formed by residues in positions 2276, 2279, 2280, and 2283, a threonine in position 1074 inserts into a pocket formed by 2272, 2275, and 2276, and a tryptophan at position 2272 contacts a leucine at position 1076. The NH_2_ of a conserved asparagine at position 2279 usually forms a hydrogen bond with the backbone carbonyl of the residue at position 1073. This interface is distinct from the DH/KR interfaces of the porcine FAS and the lovastatin PKS but most similar to the porcine FAS. The DH and KR domains of MAS do not share an interface.

While AlphaFold 3 performed exceedingly well to predict common architectures of the KS-ACP regions for 39 *Dd*PKSs, it has limits. Some, but not all, of the structures for Pks37 (also referred to as StlB) show its ER dimer on the opposite side of the KS dimer from the DH dimer. Perhaps the multiple sequence alignment used by AlphaFold 3 needs to contain more Pks37 homologs for more confident prediction. AlphaFold 3 seems not to be able to place the offloading enzymes present downstream of ACP in 21 of the *Dd*PKSs. Two of these (from Pks1 and Pks37) are dimeric chalcone synthases (PDB: 2H84) (34) likely located on the two-fold axis of the synthase, whereas the others are most likely monomeric thioester reductases (TRs). In hexameric FASs, the ACP domain is constrained by anchored, N- and C-terminal linkers. Thus, the C-terminal offloading enzymes of the *Dd*PKSs might be expected to share an interface with one of the PKS domains. However, for AlphaFold 3 to predict these putative interfaces may also require more sequence information from homologs.

### Elucidating the reaction chambers of the DdPKSs

The region within which the ACP domain diffuses to dock each of its cognate enzymes is known as the reaction chamber. The best characterized reaction chamber is that of the hexameric *Saccharomyces cerevisiae* FAS, which has been observed through cryo-EM with ACP bound to each of its cognate enzymes (42). This ACP only needs to rotate and translate within a small volume to complete a reaction cycle since its docking sites are oriented towards the center of the reaction chamber.

While it should be possible to push cryo-EM studies of Pks16 to experimentally characterize the reaction chamber of a dimeric synthase, AlphaFold 3 is already able to make atomic-level docking predictions with high confidence. It predicted how ACPs dock with downstream KSs in modular PKSs, as corroborated by mutagenesis and cryo-EM studies (43, 44). It also predicted how ACPs dock with KRs (45), DHs (46) and ERs (26) in modular PKSs, as corroborated by mutagenesis studies. In the AlphaFold 3-predicted structures of the KS-ACP regions of the 40 *Dd*PKSs, ACP domains are frequently associated with cognate enzymes. Remarkably, these interfaces are similar to those experimentally observed or predicted for modular PKSs.

As some associations dominated others, we tested various strategies to obtain more examples of less frequently observed associations. For example, because ACP/KS associations are common in the KS+DH+MT+KR_s_+ER+KR_c_+ACP dimers, 3 KS residues at the predicted ACP/KS interface (positions 86-88) were changed to aspartates. Abrogating the ACP/KS interface rather than the KS domain was preferred since KS and DH share a large interface. ACP/ER associations dominated the resulting KS*+DH+MT+KR_s_+ER+KR_c_+ACP dimers. To obtain more structures with ACP/MT and ACP/KR associations, we asked AlphaFold 3 to predict monomeric MT+KR+ACP structures.

One of the most frequent associations observed in predictions of KS+DH+MT+KR_s_+ER+KR_c_+ACP dimers is between the KS and ACP domains (**Figure 5A**). To more completely investigate these interactions, AlphaFold 3 was asked to predict how the KS+AT dimer associates with 2 copies of ACP in each of the 40 *Dd*PKSs (KSATACP_PksXX Data Files). Not only were each of the predicted structures similar to one another, they were similar to both the cryo-EM structures reported from modular PKSs in which KS associates with a downstream ACP during the extension reaction (PDBs: 7M7F, 7S6C) (23, 24). While the KSs of modular PKSs possess a second ACP docking site employed during the transacylation reaction, it is thought that a single ACP docking site is employed in both the extension and transacylation reactions conducted by iterative synthases (43, 44). Thus, ACP visits this site at least twice per the reaction cycle, once to collect an acyl chain bound to the KS reactive cysteine through the extension reaction and once to transfer the extended and processed chain back to the KS reactive cysteine. Similar to the KS/ACP interface of modular PKSs, KS makes contact with the phosphopantetheinylated end of ACP at positions 81 (usually leucine) and 84-88. Unlike the interface of modular PKSs, a loop connecting FSD and AT (positions 534-541) also makes contact with ACP residues (downstream of helix I at positions 2528-2531; this interface is similar but different for Pks1 and Pks37). In Pks2, Pks22, Pks23, Pks44, and Pks45, AT residues also make contact with ACP residues (in helices I and IV).

**Figure 5.**
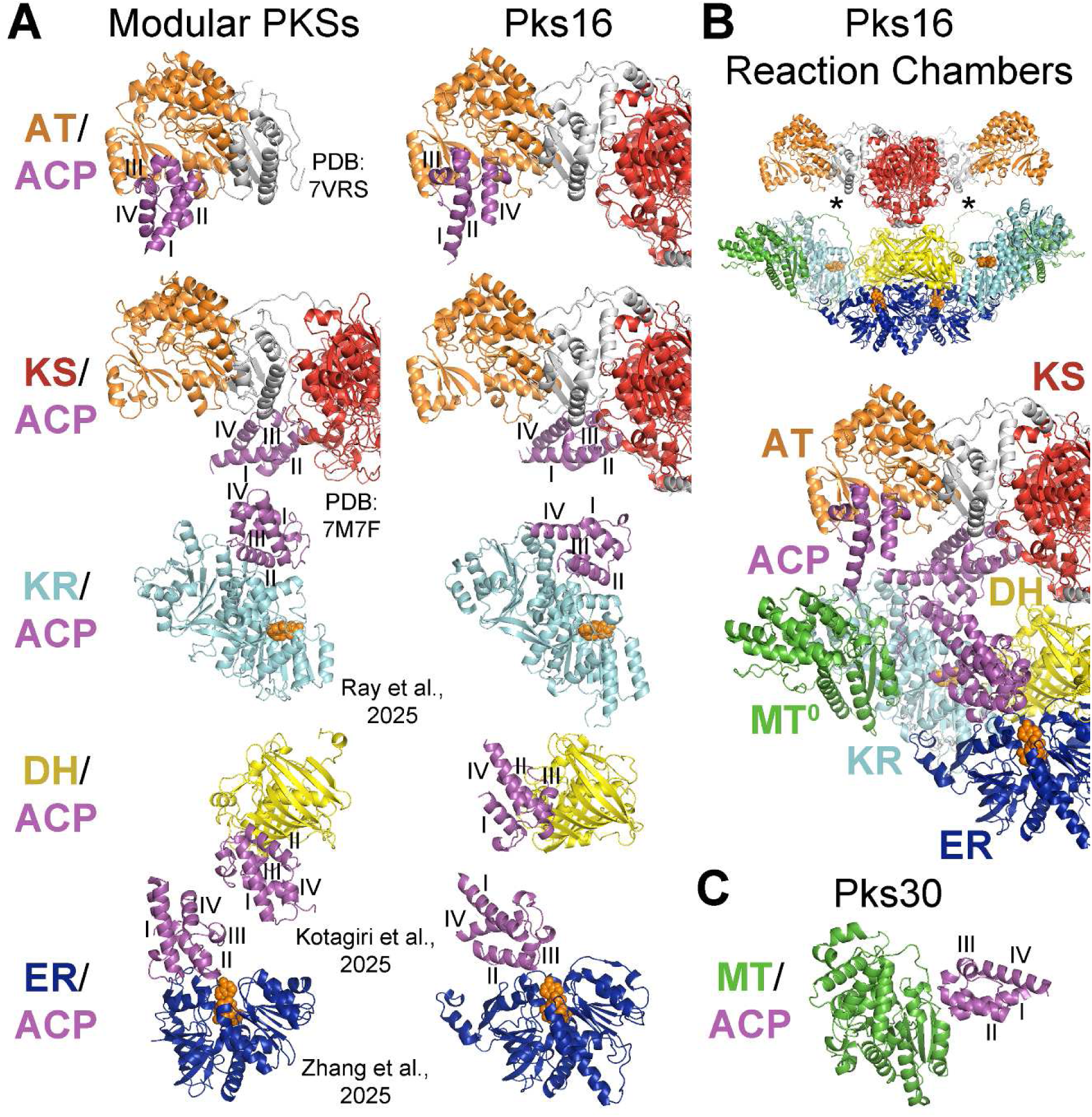
ACP docking sites predicted by AlphaFold 3 define the Pks16 reaction chamber. (A) The associations predicted for Pks16 were similar to the consensus associations for the 39 other *Dd*PKSs (the DH/ACP association is from Pks17, 96% identical to Pks16). They are similar to associations characterized in modular PKSs, the biggest exception being DH/ACP, where ACP cannot equivalently dock due to a steric clash that would result with ER. ACP helices I-IV are labeled. (B) Each Pks16 homodimer contains 2 equivalent reaction chambers (*). An overlay of docking poses maps out the reaction chamber of Pks16 and the trajectory its ACP follows during a reaction cycle (see **Movie 1**). (C) An association predicted for Pks30 is representative of the MT/ACP consensus association observed for *Dd*PKSs.

The interface observed between AT and ACP in a modular PKS (PDB: 7VRS) is relatively small (25). Perhaps the phosphopantetheinyl arm significantly contributes to the AT/ACP binding interaction, as this moiety comprises the majority of the other AT substrate, malonyl-CoA. When AlphaFold 3 was asked to predict how ATs of the *Dd*PKSs associate with ACPs, consensus solutions similar to the modular PKS AT/ACP structure were obtained for 18 PKSs (ATACP_PksXX Data Files). The residues in AT positions 733 (usually phenylalanine) and 735 generally insert into pockets formed by residues in ACP positions 2543 (the phosphopantetheinylated serine), 2547, 2550, and 2564. An AT residue at position 835 (usually phenylalanine) generally inserts into a pocket between ACP helices I and II.

While only 3 PKSs showed an MT/ACP association in the KS*+DH+MT+KR_s_+ER+KR_c_+ACP predictions, 16 more PKSs showed this consensus association in the MT+KR+ACP predictions (MTACP_PksXX.pdb Data Files). To our knowledge, MT/ACP interfaces for iterative type I PKSs have neither been characterized nor proposed in the literature. The predictions show MT associating with the phosphopantetheinylated end of ACP. The N-terminal residues of an MT helix (Pks30 positions 1508-1516; Pks30 is used as a reference for MT/ACP interactions since Pks16 does not have an active MT) make contact with the C-terminal residues of ACP helices I and II (Pks30 positions 2550-2551 and 2567-2575). A conserved serine (Pks30 position 1508) forms a hydrogen bond with the DSL aspartate (Pks30 position 2569) of ACP.

Eleven PKSs showed a KR/ACP association in the KS*+DH+MT+KR_s_+ER+KR_c_+ACP predictions, and 10 more PKSs showed this consensus association in the MT+KR+ACP predictions (KRACP_PksXX.pdb Data Files). The interfaces are broadly similar to one another as well as to the interface predicted for B-type KRs from modular PKSs (45). The location of the phosphopantetheinyl arm attached to the conserved ACP serine (position 2543) affects how polyketides enter the KR active site and thus the stereochemical outcome of the reduction reaction. The predictions show how interactions between residues in ACP helices II and III (positions 2542-2569) and residues near the KR active site (positions 1782-1784, 2335, 2385-2386, 2428-2438, and 2478-2481) dictate the placement of the phosphopantetheinyl arm. Generally, the greater interaction KR residues with ACP helix III residues leads to a rotation of ACP relative to how it is bound in modular PKSs.

The associations of DH and ER with ACP are apparently interconnected, with ER residues aiding the DH/ACP association and DH residues aiding the ER/ACP association. DH may be more reliant on ER, as 4 PKSs in the KS*+DH+MT+KR_s_+ER+KR_c_+ACP predictions show the DH/ACP consensus association compared to 27 PKSs that show the ER/ACP consensus association (DHACP_PksXX.pdb and ERACP_PksXX.pdb Data Files). When AlphaFold 3 is asked to predict DH/ACP associations in the absence of other domains, no consensus docking is observed. The DH and ACP domains cannot associate as predicted for modular PKSs due to steric clashes with the ER dimer (ERs in modular PKS are almost exclusively monomeric) (46, 47). In the consensus association observed for the *Dd*PKSs, ACP is relatively rotated 180°, similar to how AlphaFold 3 predicts ACP docks with DH in the porcine FAS and MAS. The ER residues that aid in the DH/ACP association (ER residues in positions 1857, 1890-1893, and 2175 contact ACP residues in positions 2524-2539, C-terminal to helix I) also aid in the ER/ACP association. Residues close to the DH active site (positions 1003-1004, 1043, 1149, and 1210) only make a few contacts with residues near the phosphopantetheinylated serine (position 2543) on ACP helices II and III.

The ER/ACP associations observed from 27 PKSs in the KS*+DH+MT+KR_s_+ER+KR_c_+ACP predictions are consistent. ER residues (positions 1856-1857, 1859, 1890-1893, 2171, and 2175) make contact with ACP residues in and between helices II and III (positions 2543-2565). At the center of these interactions is contact between 2 residues at the C-terminal end of an ER helix (positions 1890-1891) and an ACP side chain at position 2564 (predominantly isoleucine). These ER residues usually also make contact with an ACP valine conserved in position 2547. The side chains of ER residues at positions 1857, 1859, and 2171 (most often asparagine, glutamate, and lysine, respectively) usually form a hydrogen bond network with an ACP asparagine conserved in position 2560. A β-hairpin motif in DH [DxK(S/T)NEWI, not present in Pks16 or Pks17, but comprised of residues equivalent to those at positions 1035-1043] and neighboring DH residues make highly complementary interactions with residues between ACP helices I and II (positions 2529-2538). These contacts include a hydrogen bond between the aspartate from the DH β-hairpin motif and a conserved ACP asparagine (position 2531), a hydrophobic interaction between the isoleucine of the DH β-hairpin motif and a conserved ACP leucine (position 2534), a hydrogen bond between a DH serine/threonine with a semi-conserved ACP aspartate (position 2538), and a salt bridge between a DH lysine and a conserved ACP aspartate (position 2533). The ACP/ER association is similar to, but distinct from, the association AlphaFold 3 predicts for the porcine FAS, MAS, and modular PKSs (26).

From the KS/ACP, AT/ACP, KR/ACP, DH/ACP, and ER/ACP associations predicted for Pks16, a movie was generated showing how ACP moves during a reaction cycle in the synthesis of a fatty acid (**Movie 1**). ACP does not need to translate much in its reaction chamber (**Figure 5B**). The reaction chambers of the other *Dd*PKSs may slightly differ due to the presence of C-terminal domains that further constrain ACP movement or functional MTs with which ACP must associate (**Figure 5C**). However, given the structural relationships between Pks16 and other dimeric FASs/PKSs in amoeba, mammals, fungi, and bacteria, these molecular machines are likely to operate through similar dynamics.

## Conclusions

In this pilot study, we employed Shotgun-EM to analyze *D. discoideum* cell lysates, successfully isolating several of its megadalton multiprotein assemblies. By employing this methodology with heterogeneous mixtures, we were able to identify and reconstruct several protein complexes, including an octahedral assembly of the dihydrolipoyllysine-residue succinyltransferase subunit of the mitochondrial 2-oxoglutarate dehydrogenase complex, the 20S proteasome, and an unidentified hexameric star-like protein complex. Most notably, we serendipitously obtained a 3.9 Å experimental structure for a *D. discoideum* polyketide synthase, Pks16, the first of 40 *D. discoideum* PKSs to have an experimentally determined structure, which allowed us to perform a comparative analysis across the members of this important protein family and computationally characterize the reaction chambers of this important class of proteins.

## Abbreviations

Cryo-EM: cryogenic electron microscopy
PKS: polyketide synthase; size-exclusion chromatography
IEX: ion exchange chromatography
MS: mass spectrometry
ACP: acyl-carrier protein
AT: acyltransferase
KS: ketosynthase
MT: methyltransferase
SAM: *S*-adenosylmethionine
KR: ketoreductase
DH: dehydratase
ER: enoylreductase

## Acknowledgments

This research was funded by grants from the National Institutes of Health (R01 GM106112 to A.T.K.-C., R35 GM122480 to E.M.M.), Army Research Office (W911NF-12-1-0390 to E.M.M.), and the Welch Foundation (F-1712 to A.T.K.-C., F-1515 to E.M.M.). The authors thank Shane Gonen for the suggestion of testing HsLV and Simone Ritchey for assistance with culturing *D. discoideum*.

## Authors’ contributions

Gabriel Hoogerbrugge: Conceptualization; Formal analysis; Investigation; Visualization; Methodology; Writing—original draft; Writing—review and editing. Adrian T Keatinge-Clay: Conceptualization; Formal analysis; Investigation; Visualization; Resources; Methodology; Writing—original draft; Writing—review and editing. Edward M Marcotte: Conceptualization; Formal analysis; Investigation; Visualization; Resources; Supervision; Methodology; Writing—original draft; Writing—review and editing.

## Data availability

Mass spectrometry proteomics data was deposited in the MassIVE/ProteomeXchange database (48) under accession number PXD061189/MSV000097213 (doi:10.25345/C5S46HJ6K). Cryo-electron microscopy data was deposited in the Electron Microscopy Data Bank (49) under accession numbers EMD-49490 (Odo2), EMD-49491 (Pks16), EMD-49492 (Hexameric star complex), EMD-49493(20S proteasome). Coordinates for Pks16 and Odo2 were deposited in the Protein Data Bank as PDB 9NJU and 9NJT, respectively. Coordinates for all computationally predicted structures, including the 40 PKS proteins and modeled subsets of domains, are available through a Zenodo repository (10.5281/zenodo.14925470).

## In Brief statement

Large protein complexes drive essential cellular processes, yet many remain structurally uncharacterized. By combining native chromatography, cryo-electron microscopy, and mass spectrometry, we analyzed a sampling of endogenous megadalton protein assemblies from the slime mold *Dictyostelium discoideum*. This study uncovered multiple protein structures, including for the polyketide synthase Pks16, the primary fatty acid synthase in slime molds, allowing us to perform a comparative analysis across *Dictyostelium’s* full repertoire of >40 PKS proteins.

## Accompanying files

### Supplemental figures

**Figure S1-4: Cryo-EM workflow and statistics for each complex**

**Figure S5: Multiple sequence alignment of *Dd*PKSs**

## Supplemental tables

**Table S1: Mass spectrometry protein identifications**

**Table S2: Cryo-EM statistics**

