## Supplemental Figures and Legends for "Serendipity and the slime mold: a visual survey of megadalton protein assemblies reveals the structure of the polyketide synthase Pks16"

**Supplemental Figure Legends**

**Figures S1-4. Cryo-EM processing pipelines for (S1) polyketide synthase Pks16, (S2) the octahedral Odo2 dihydrolipoyllysine-residue succinyltransferase complex, (S3) the 20S proteasome, and (S4) the hexameric star complex.** In each case, the specific cryo-EM workflow is presented with a view of the final EM density along with the statistics of the final reconstruction. (A) Data processing and refinement pipeline for the complex using cryoSPARC v4.5. The symmetry utilized at each refinement step is noted at the bottom right hand corner of the respective refinement. (B) Cryo-EM volume colored by local resolution estimation (units in Å). (C) Gold-standard Fourier shell correlation (GSFSC) curve using an FSC threshold of 0.143, as calculated by cryoSPARC. (D) Viewing angle distribution plot.

**Figure S5. Multiple sequence alignment for the KS-ACP regions of the *Dictyostelium discoideum* PKSs (except Pks37).** Domain boundaries are indicated with colored bars. Numbering and secondary structure is based on Pks16.

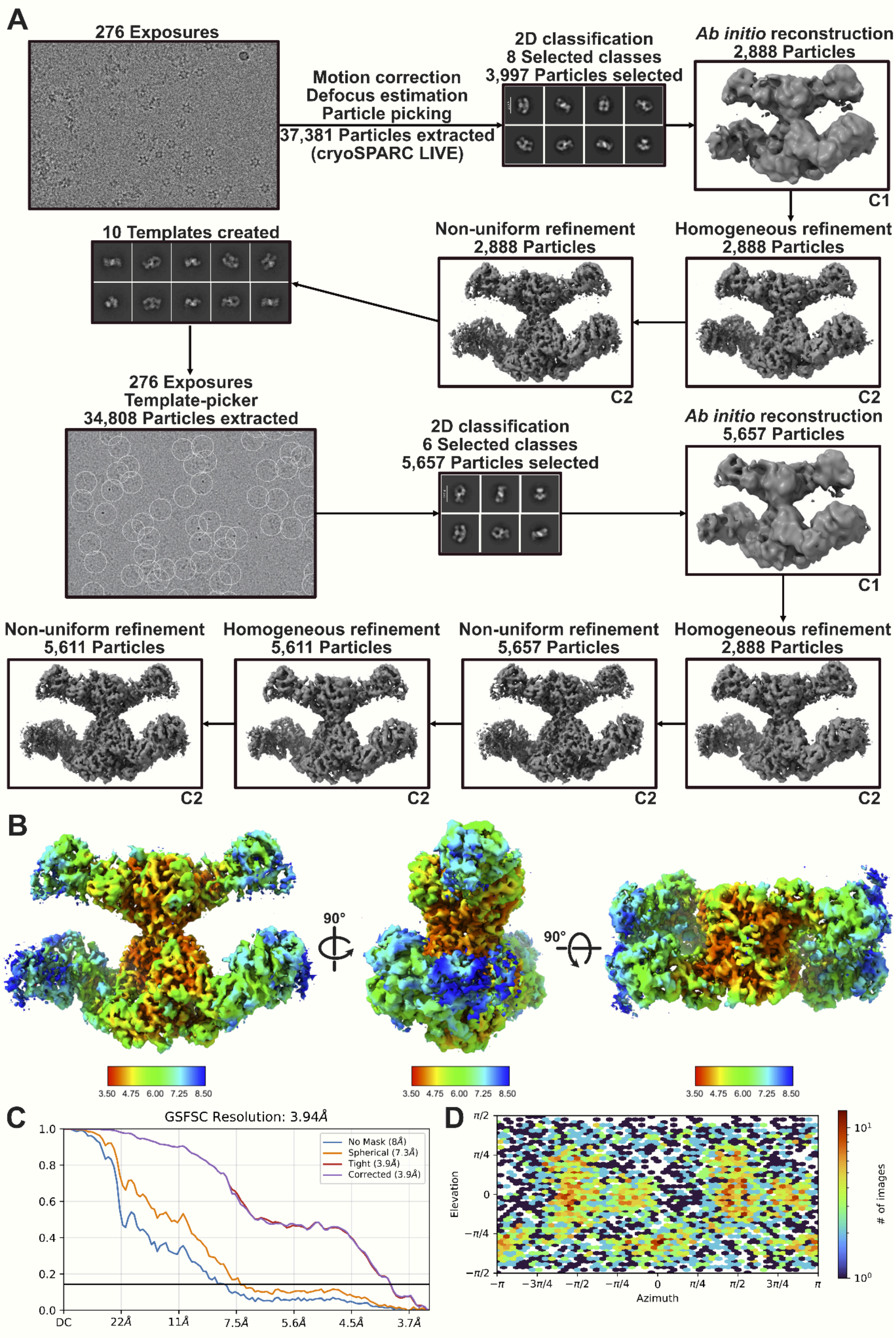


**Figure S1. Cryo-EM processing pipeline for the *D. discoideum* polyketide synthase Pks16.**

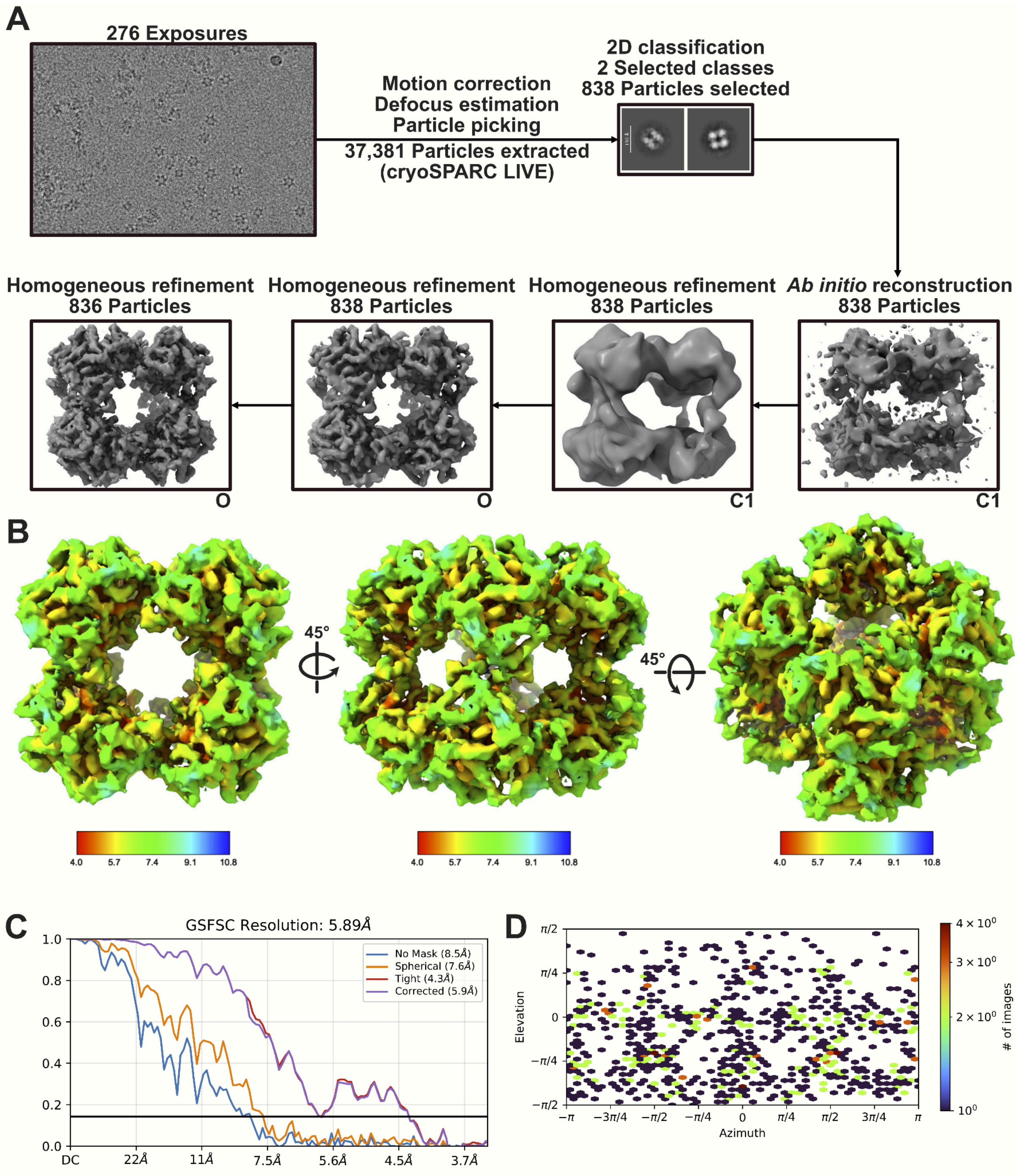


**Figure S2. Cryo-EM processing pipeline for the octohedral assembly of the *D. discoideum* Odo2 dihydrolipoyllysine-residue succinyltransferase.**

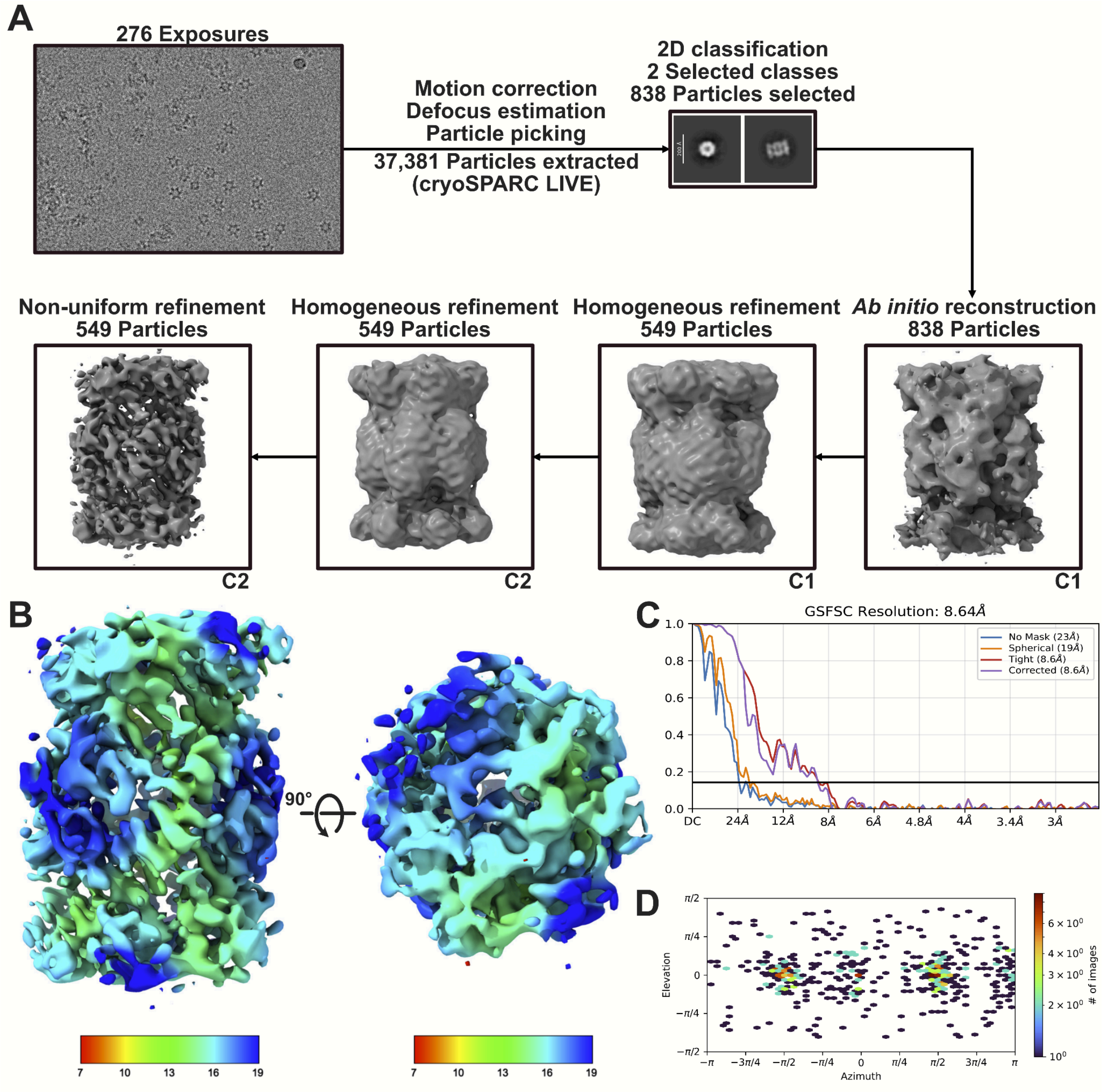


**Figure S3. Cryo-EM processing pipeline for the *D. discoideum* 20S proteasome.**

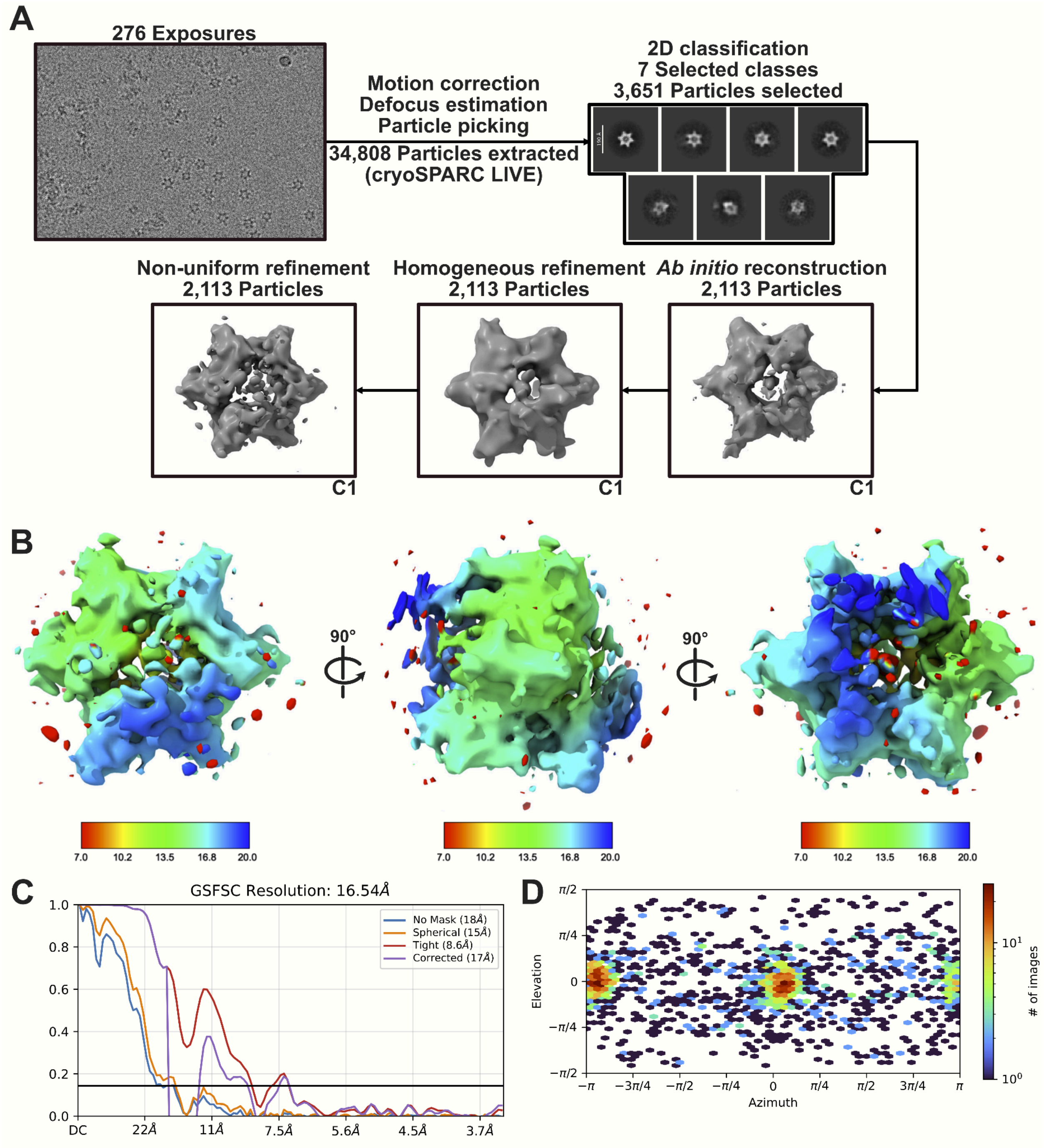


**Figure S4. Cryo-EM processing pipeline for the *D. discoideum* hexameric star complex.**

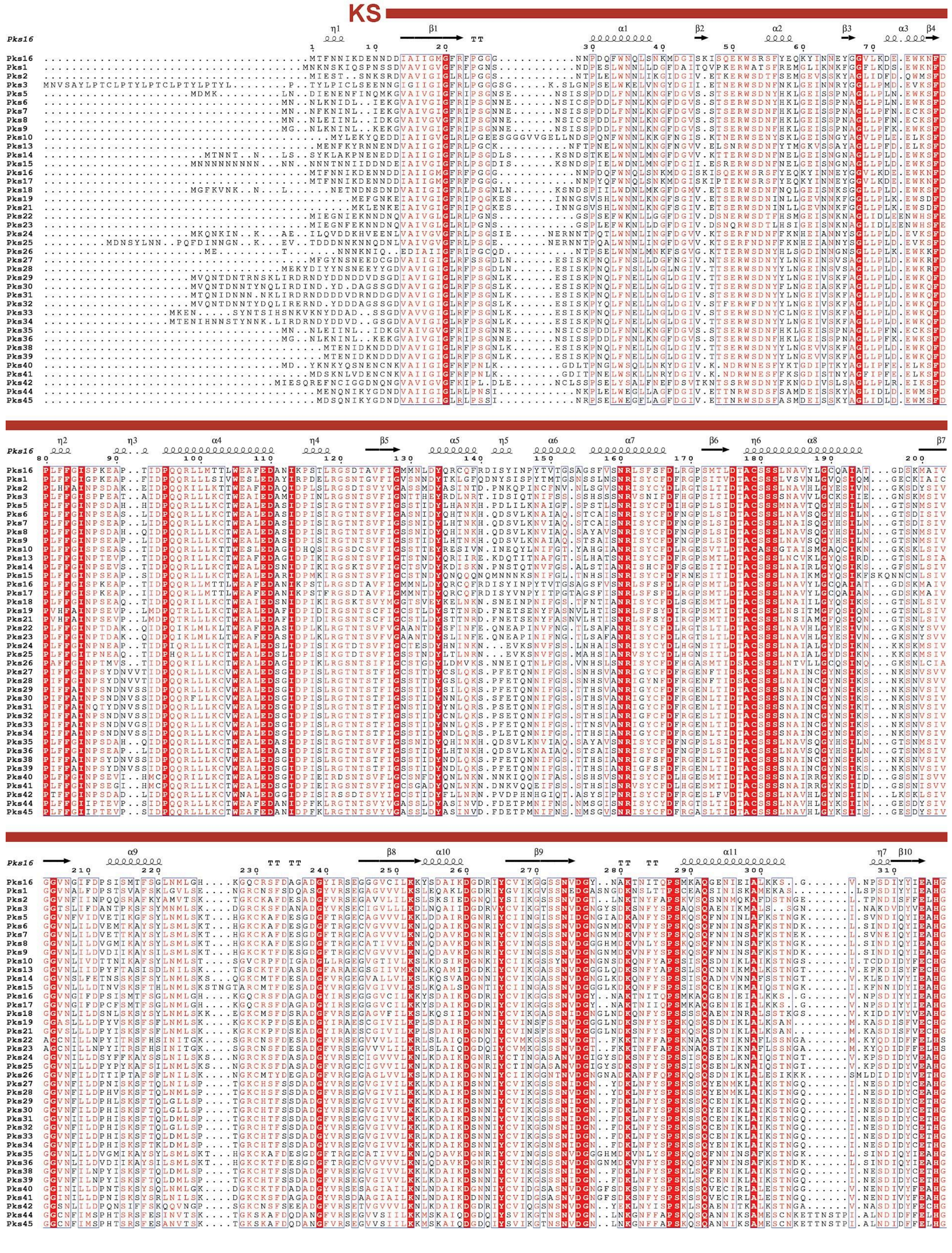
KS

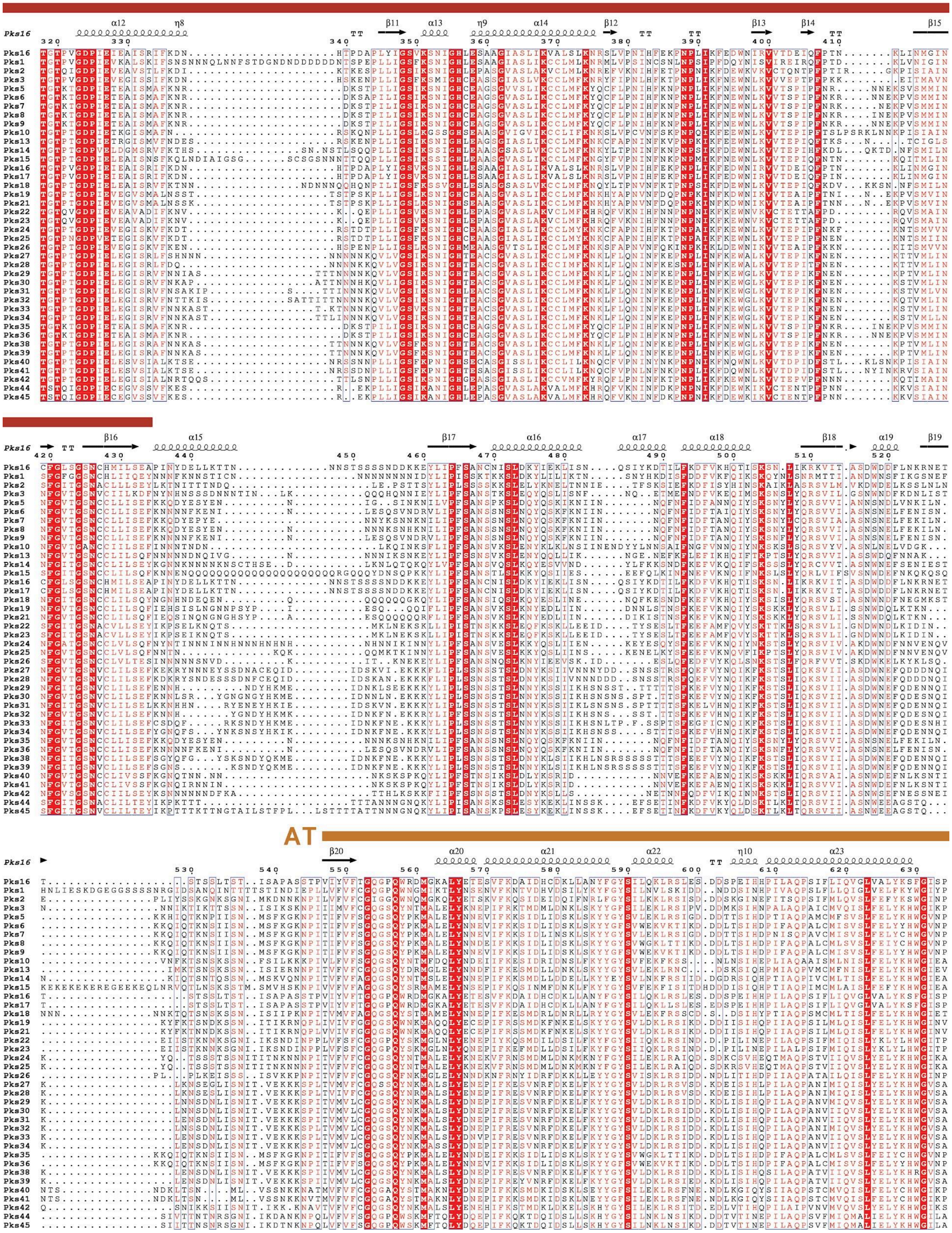

AT

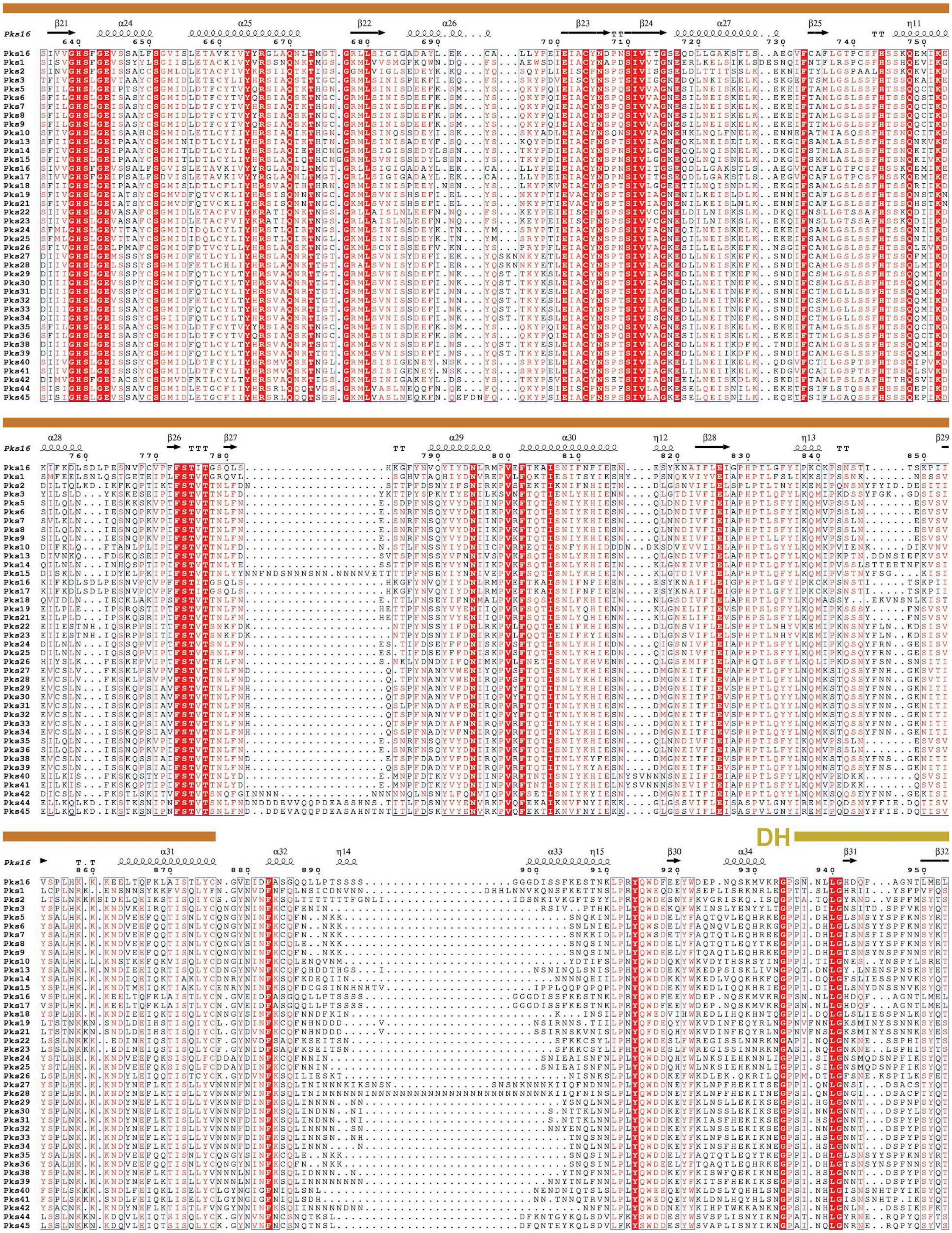
DH

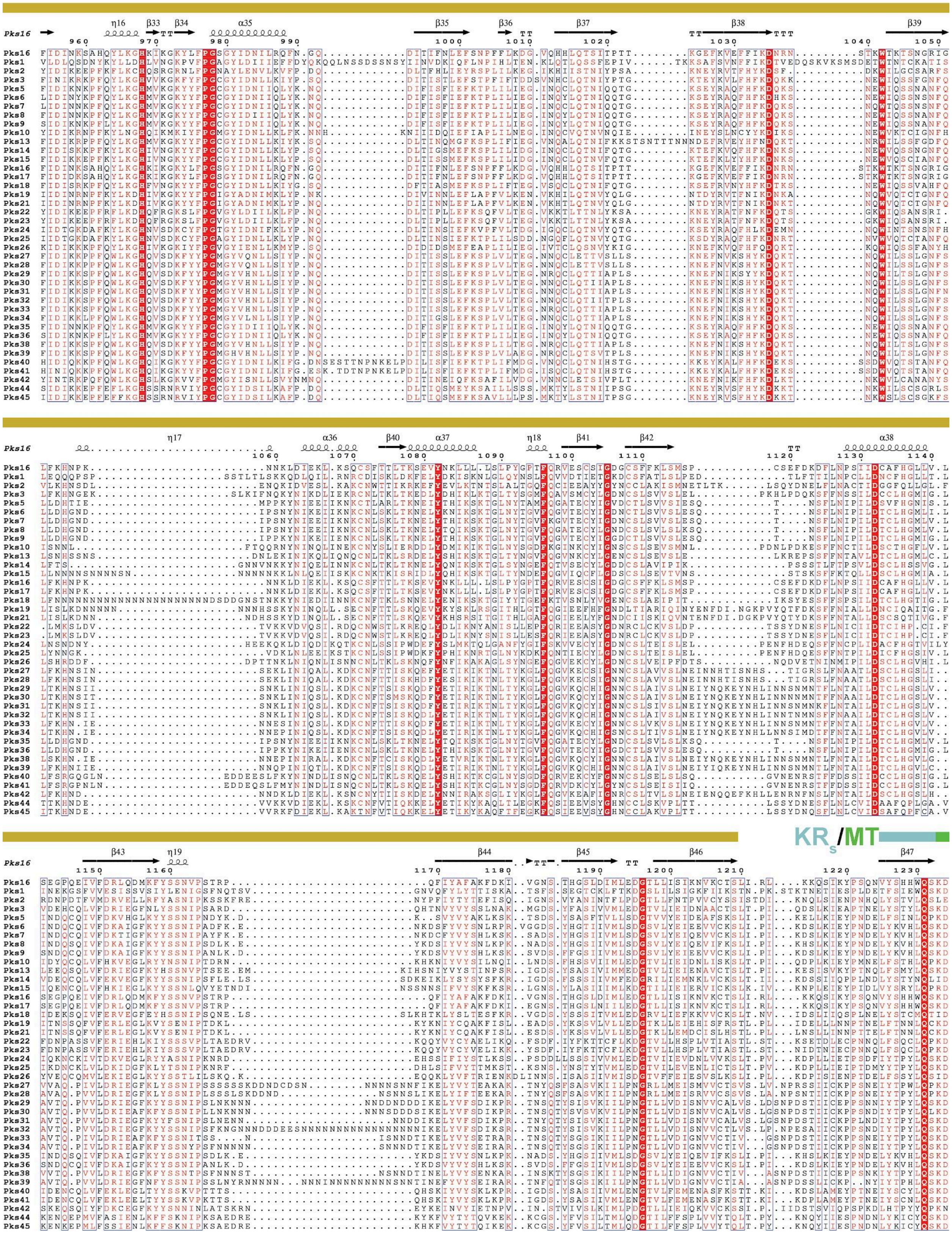
KRs/MT

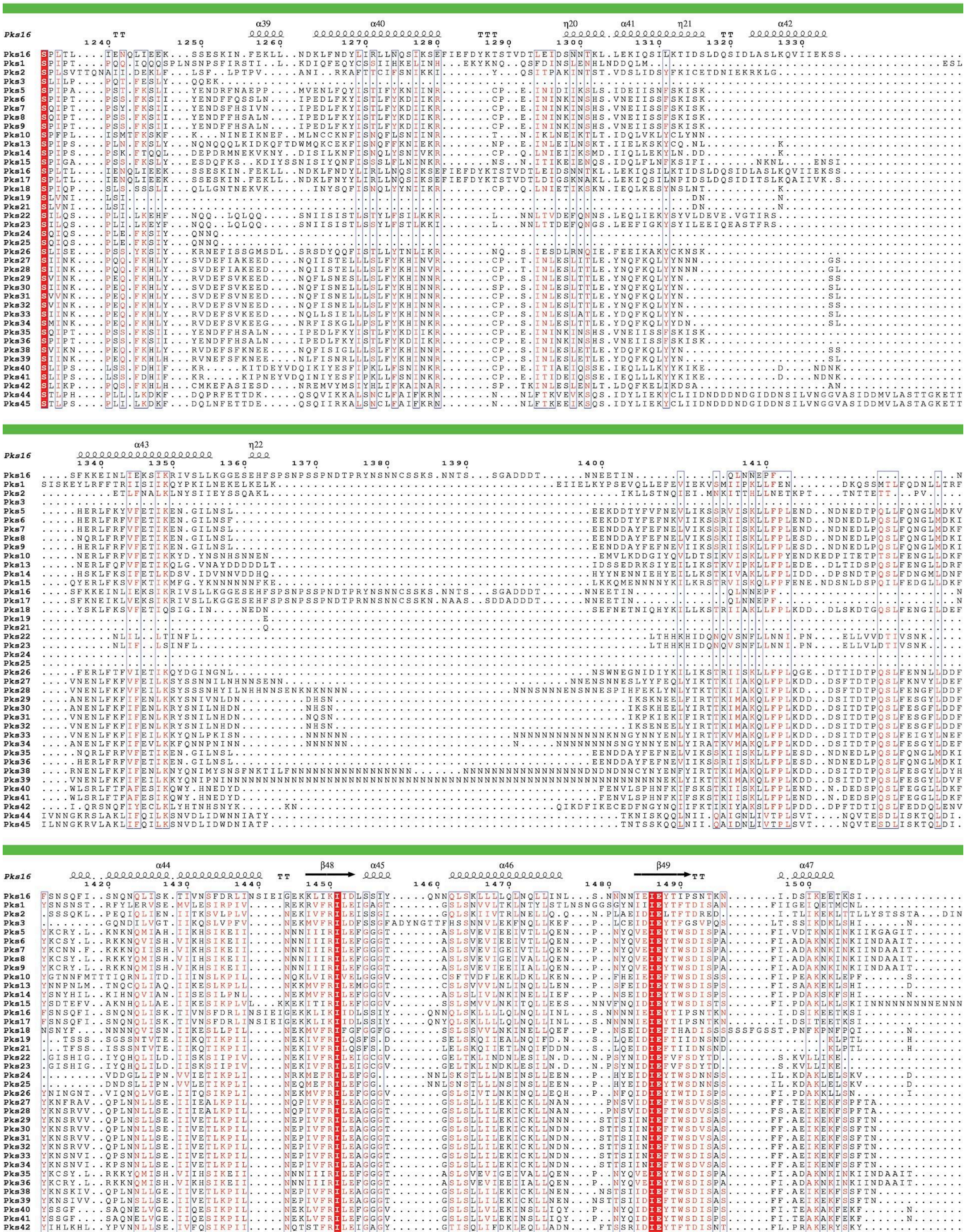

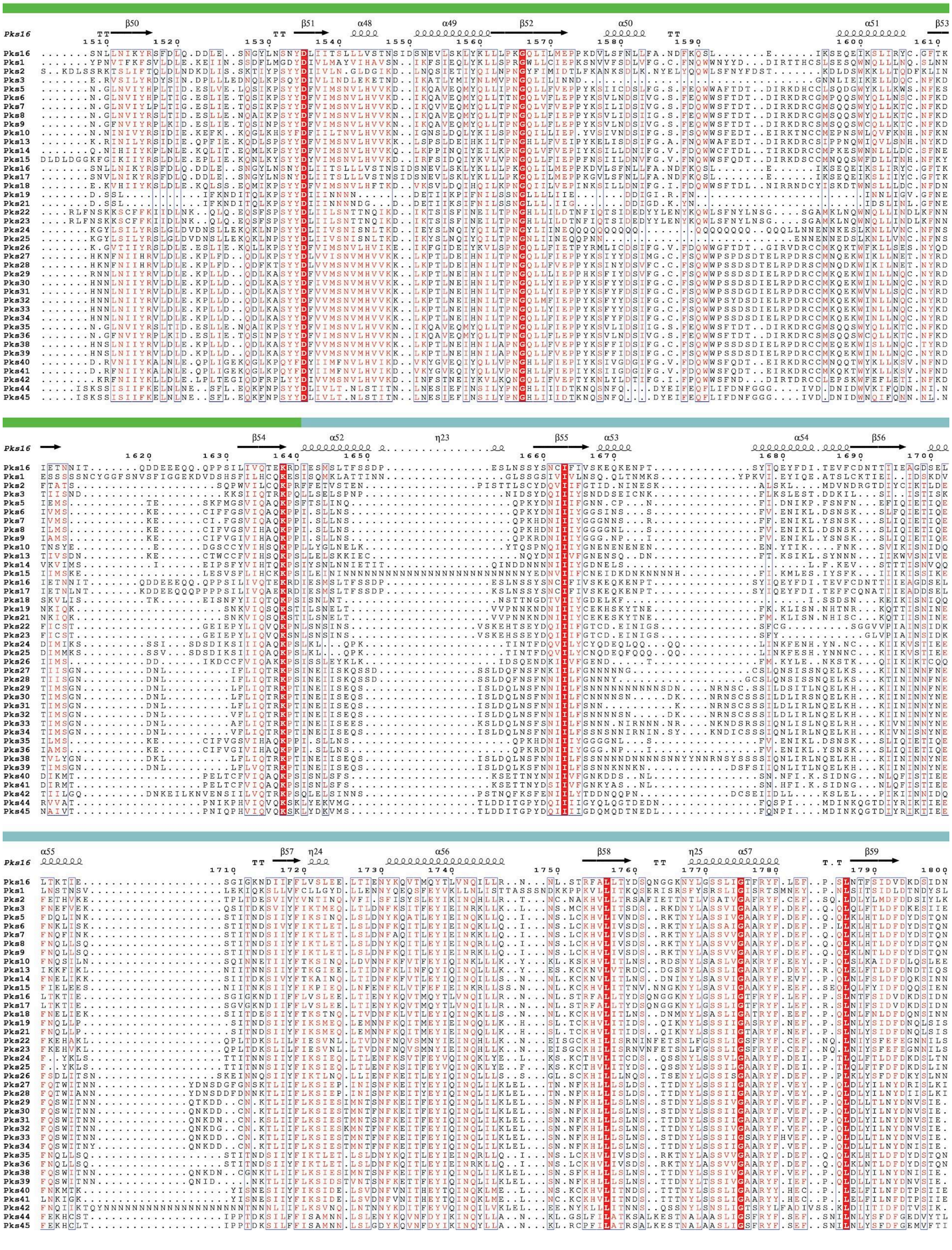
Pks16
β50 β51 α48 α49 β52 α50 α51 β53
β54 α52 η23 β55 α53 α54 β56
α55 β57 η24 α56 β58 η25 α57 β59
Pks16
Pks1
Pks2
Pks3
Pks5
Pks6
Pks7
Pks8
Pks9
Pks10
Pks13
Pks14
Pks15
Pks16
Pks17
Pks18
Pks19
Pks21
Pks22
Pks23
Pks24
Pks25
Pks26
Pks27
Pks28
Pks29
Pks30
Pks31
Pks32
Pks33
Pks34
Pks35
Pks36
Pks38
Pks39
Pks40
Pks41
Pks42
Pks44
Pks45

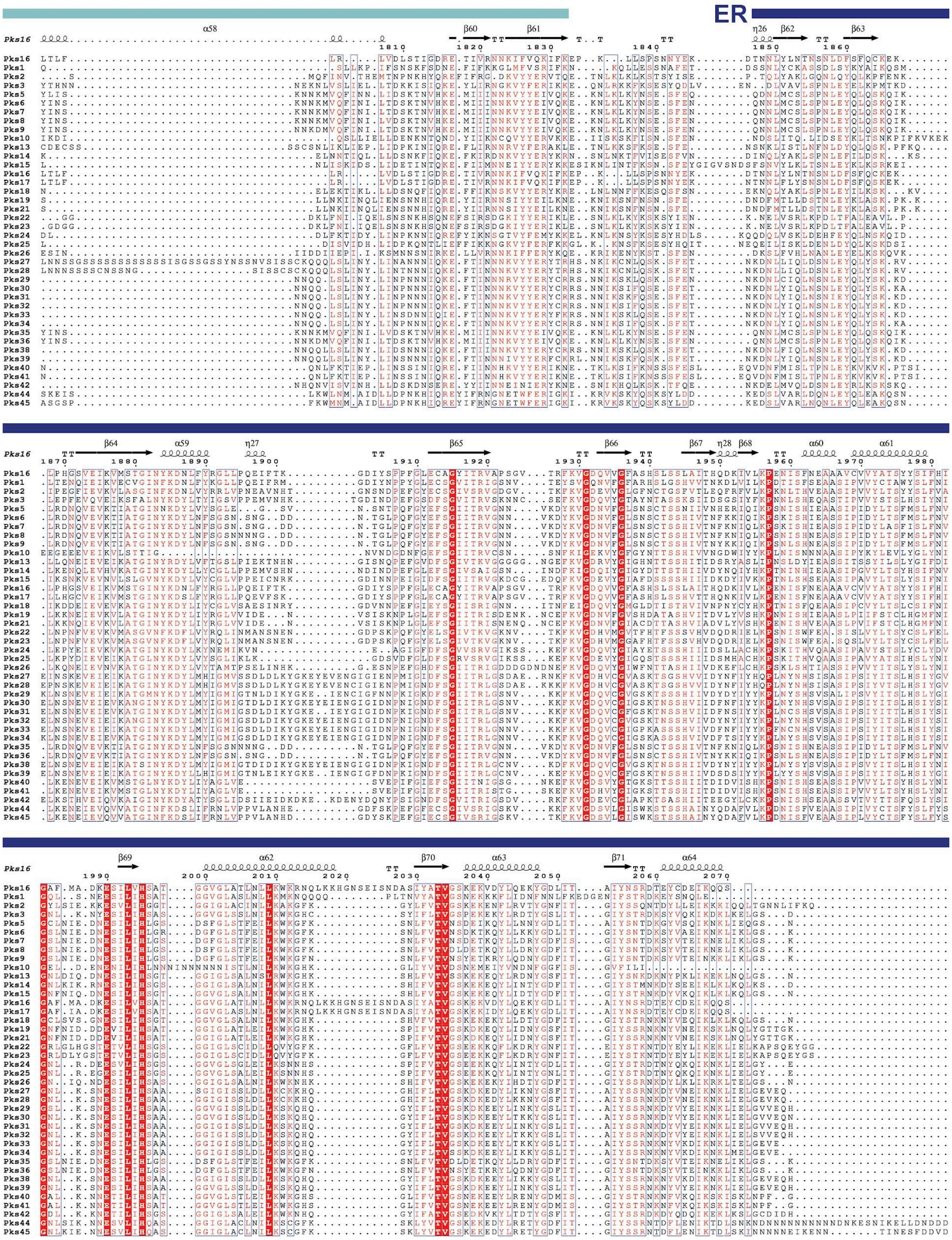
ER

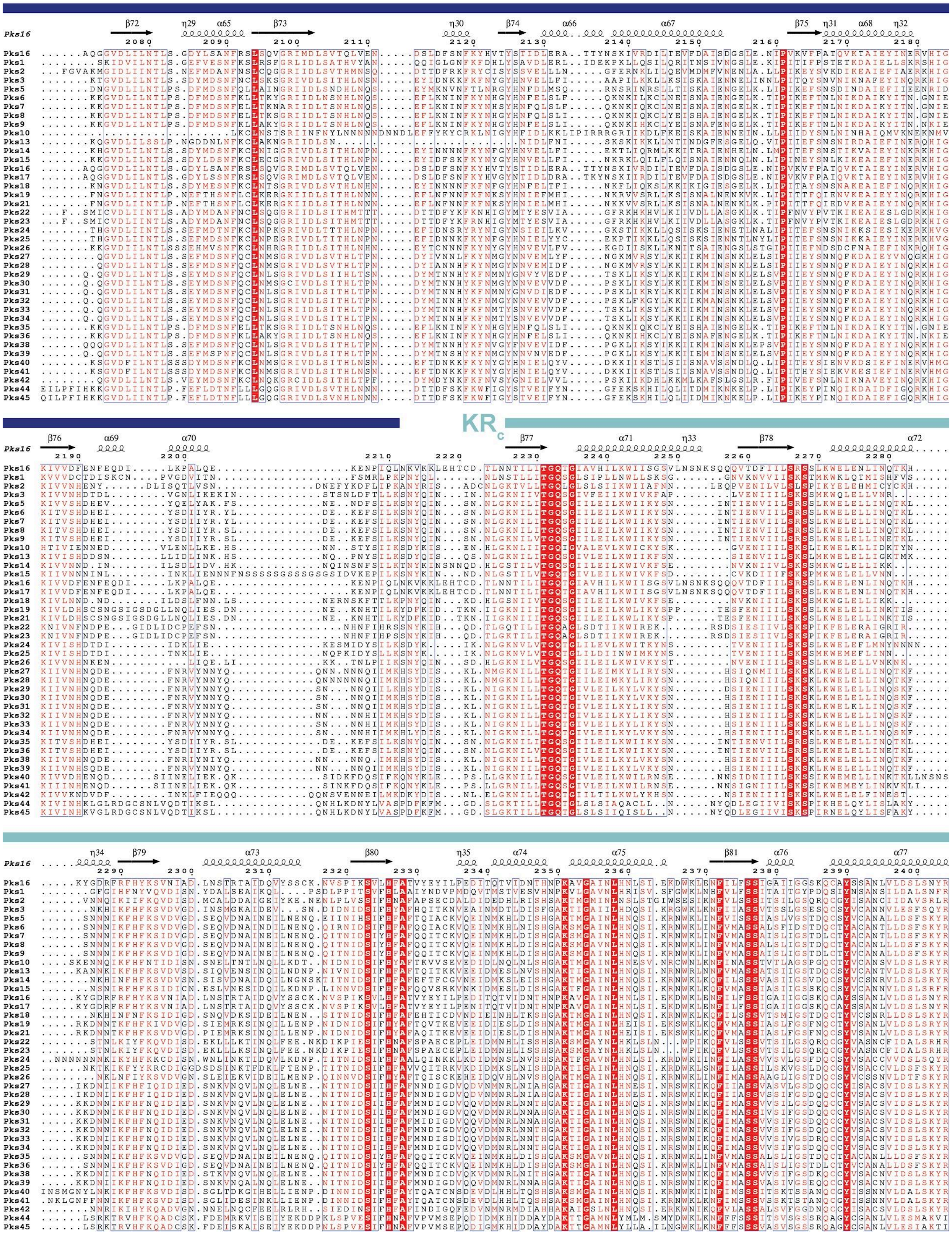
KRc
Pks16

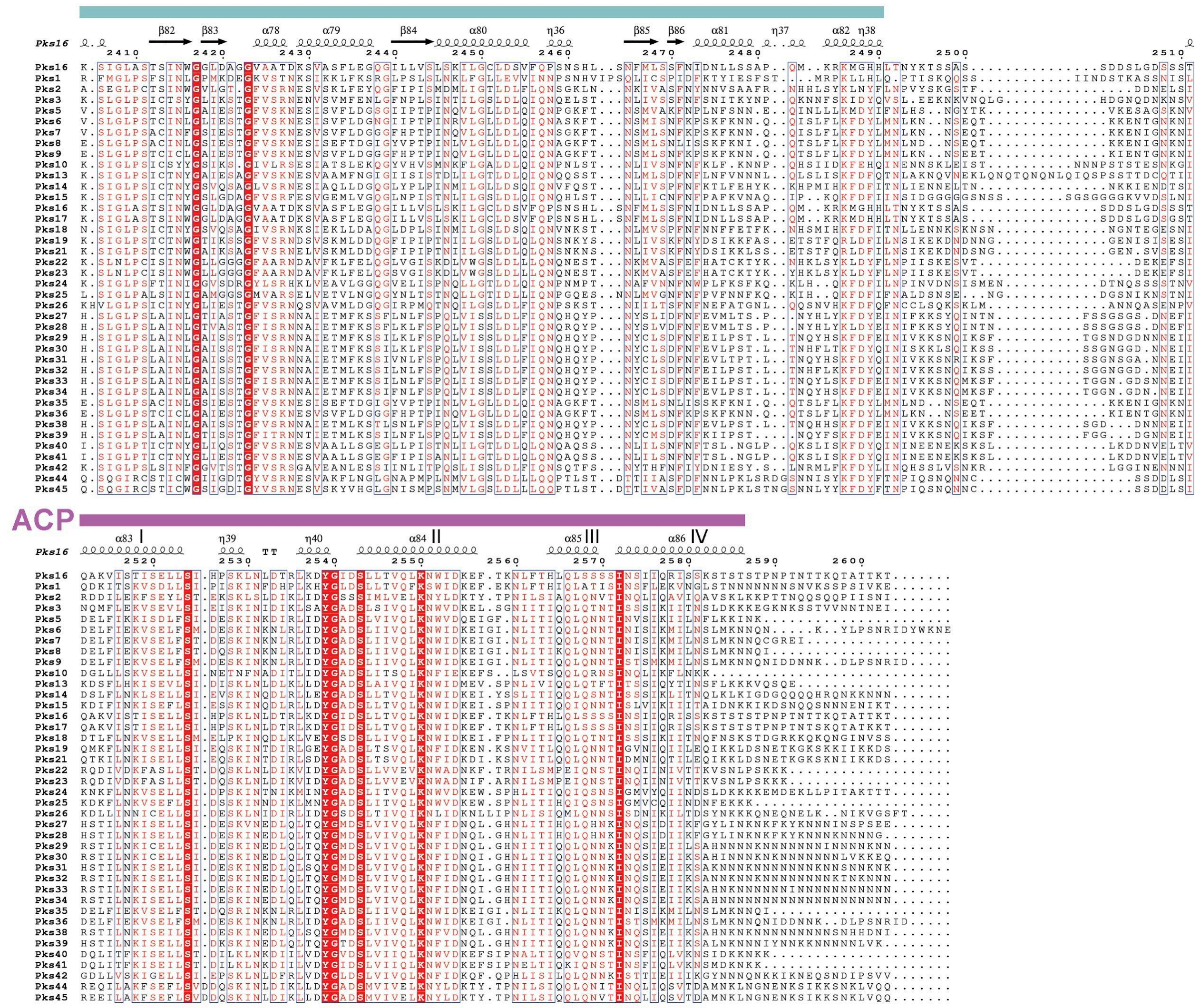


**Figure S5. Multiple sequence alignment for the KS-ACP regions of the *Dictyostelium discoideum* PKSs (except Pks37).**
